# Arf GAP containing dAsap regulates NMJ organization and synaptic calcium through Arf6-dependent signaling in *Drosophila*

**DOI:** 10.1101/2023.06.30.547304

**Authors:** Bhagaban Mallik, Shikha Kushwaha, Anjali Bisht, MJ Harsha, C. Andrew Frank, Vimlesh Kumar

## Abstract

Synaptic morphogenesis involves an interplay of multiple signaling pathways and requires membrane remodeling and cytoskeleton dynamics. We identified the BAR-domain protein dAsap (Arf GAP, SH3, Ankyrin repeat, and PH domain) as one of the regulators of synaptic morphogenesis. Loss of dAsap results in decreased bouton numbers, increased inter-bouton diameter, and disrupted microtubule organization at the nerve terminals. Electrophysiological analysis of the mutants revealed a gain in neurotransmission compared to control neuromuscular junctions (NMJs). *dAsap* mutant NMJs have increased evoked amplitude, increased spontaneous miniature frequency, and significantly fewer synaptic failures in low calcium. Consistent with these observations, *dAsap* mutants have increased active zone number. Additional pharmacological and genetic manipulations that are known to impair calcium release from stores suppress the dAsap phenotypes. Finally, we show that expressing a GDP-locked form of Arf6 in *dAsap* mutants restored the NMJ morphological defects, disrupted cytoskeleton, and aberrant neurotransmission. Thus, we propose a model in which dAsap regulates NMJ morphogenesis and synaptic calcium homeostasis through Arf6-dependent neuronal signaling.

## Introduction

Synapse formation, organization, and remodeling are regulated by membrane-binding proteins that interact with the underlying actin/microtubule-based cytoskeleton. Cell culture and genetic studies have highlighted the role of Bin-Amphiphysin-Rvs (BAR) domain superfamily proteins as potential regulators of synapse remodeling and physiology (Mallik et al., 2022; Mallik et al., 2017; Miura et al., 2016; Stanishneva-Konovalova et al., 2016; Ukken et al., 2016). Many BAR-domain-containing proteins are multidomain proteins and possess modules that may modulate neuronal signaling by regulating the activities of small GTPase (Nadif Kasri et al., 2009; Szatmari et al., 2013). For instance, BAR-domain proteins like GTPase Regulator Associated with Focal Adhesion Kinase (GRAF) and Oligophrenin-1 contain GAP domains that regulate RhoA-dependent cytoskeleton dynamics (Billuart et al., 1998; Taylor et al., 1999). Other BAR-domain proteins like ASAP and CentaurinB1A contain the ArfGAP domain that regulates Arf-dependent cellular signaling (Moore et al., 2007; Rodrigues et al., 2016; Szatmari et al., 2021).

Arfs regulate diverse cellular processes such as membrane trafficking, actin remodeling, endosomal trafficking, cell migration, cancer invasion, phospholipid metabolism, calcium regulation, endocytosis, and phagocytosis (Adarska et al., 2021; D’Souza-Schorey and Chavrier, 2006; Ha et al., 2008b; Ismail et al., 2010; Nacke et al., 2021; Pelletan et al., 2015; Randazzo et al., 2007; Sabe, 2003; Shi and Grant, 2013; Tanna et al., 2019). While Arf1 has been shown to regulate the polarization of the Golgi stack, Arf6 facilitates the recycling of endosomes, organization of the cortical cytoskeleton, endocytosis of synaptic vesicles, development and stability of dendritic spines and regulation of axonal growth in developing neurons (Bannykh et al., 2005; Choi et al., 2006; Hernandez-Deviez et al., 2002; Hernandez-Deviez et al., 2004; Kim et al., 2015; Krauss et al., 2003; Tagliatti et al., 2016). Moreover, Arf1 and Arf6 have been shown to regulate AMPA receptor trafficking and spine formation in cultured hippocampal neurons (Miyazaki et al., 2005; Rocca et al., 2013). Thus, regulating Arfs in the neuron is crucial for synapse development and synaptic plasticity (Hernandez-Deviez et al., 2007; Kim et al., 2020).

Arfs are regulated by Arf-specific GTPase-activating proteins (ArfGAPs) which potentiate GTP hydrolysis and promote an exchange of GTP to GDP in cells. One such protein class belongs to the ASAP subfamily of BAR domain proteins. These are characterized by the presence of a conserved ArfGAP domain and stimulate hydrolysis of GTP by its intrinsic GTPase activity (Ismail *et al*., 2010). Furthermore, ASAP proteins exhibit ArfGAP activity for Arf1, Arf5, and Arf6 (Brown et al., 1998; Ismail *et al*., 2010; Johnson et al., 2011; Rodrigues *et al*., 2016). Similar to Arfs, ASAP is implicated in cell migration, invasion, podosome formation, remodeling of cytoskeletal proteins, and endosomal recycling of various cargos in cells (Bharti et al., 2007; Chen et al., 2020; Gasilina et al., 2019; Ha et al., 2008a; Inoue et al., 2008; Randazzo et al., 2000). Moreover, *in vitro* study revealed that ASAP can bind to Arf6 and stimulates calcium-mediated GTP hydrolysis (Ismail *et al*., 2010). Despite several structural and biochemical studies, ASAP-mediated Arfs function and calcium maintenance in neurons remain elusive at the organism level.

A forward genetic screen identified dAsap as one of the critical regulators of the *Drosophila* NMJs (Mallik *et al*., 2017). In order to understand the mechanism by which dAsap regulates NMJ organization, we first created loss-of-function mutations of *dAsap* using CRISPR/Cas9-based genome editing technology. While the *dAsap* mutants were viable, they showed severe defects in NMJ organization. *dAsap* mutants have increased mEJP frequency and showed significantly reduced synaptic failures under low extracellular calcium. The biochemical and epistatic interaction analysis revealed that dAsap regulates Arf6 activities. Further studies support two possible reasons for increased miniature frequency and evoked release: a) Loss of dAsap promote more calcium release from the intracellular stores, and b) increased active zone number at the nerve terminals in dAsap mutants.

Interestingly, overexpression of a dominant negative form of Arf6 rescues structural and functional defects of *dAsap* mutants. Besides, the elevated evoked activity was further suppressed to the control level by blocking inositol triphosphate receptor (IP3R) and ryanodine receptors (RyR), further suggesting endoplasmic reticulum mediated calcium release controls evoked activity at the *dAsap* mutant NMJs. Additionally, the elevated evoked release and miniature frequencies were further suppressed by chelating calcium intracellularly with membrane-permeable BAPTA-AM. These findings suggest that IP3R and RyR-mediated IP3-directed signaling might potentiate evoked response in *dAsap* mutants. These results are consistent with the model that the control of intracellular calcium dynamics is essential for neurotransmitter release. Thus, we suggest that dAsap-mediated Arf6-dependent signaling regulates *Drosophila* NMJ organization and synaptic calcium homeostasis.

## Results

### BAR domain of dAsap deforms synthetic liposomes *in vitro* but does not induce membrane tubules in cultured S2R+ cells

The *Drosophila* Asap (dAsap) is a multidomain protein consisting of a conserved N-BAR domain at its N-terminal and an SH3 domain at the C-terminal. It also contains a PH domain, an Arf GAP domain, and two ankyrin repeats (Figure 1A). Earlier reports have shown that the BAR domains bind to phospholipids and deform the synthetic membranes to generate membrane tubules and vesicles (Mallik *et al*., 2017; Masuda et al., 2006). To assess whether the BAR domain of dAsap is biochemically conserved, we purified the BAR domain of dAsap (dAsap^N-BAR^) and tested its ability to deform the synthetic liposomes. We found that the GST-tagged dAsap^N-BAR^ induced membrane tubules within 10 minutes of incubation with liposomes (Figure 1B-C). This data suggest that the dAsap^N-BAR^ domain can deform synthetic liposomes *in vitro*, similar to other N-BAR domain-containing proteins. However, overexpression of dAsap^N-BAR^ did not induce any detectable tubules or vesicles in the cultured S2R+ cells (Figure 1D, F, and H). Interestingly, overexpression of full-length dAsap (*UAS-dAsap^FL^*) or dAsap lacking the BAR domain (*UAS-dAsap^ΔBAR^*) causes massive induction of actin-rich filopodia in S2R+ cells (control, 0.38 ± 0.10; *dAsap^BAR^*, 1.56 ± 0.30; *dAsap^FL^*, 5.86 ± 0.60; *dAsap****^Δ^****^BAR^*, 5.56 ± 0.71) (Figure 1E, G and H). These data indicate that the N-BAR domain of dAsap is dispensable for membrane tubule formation in cultured cells.

**Figure 1.**
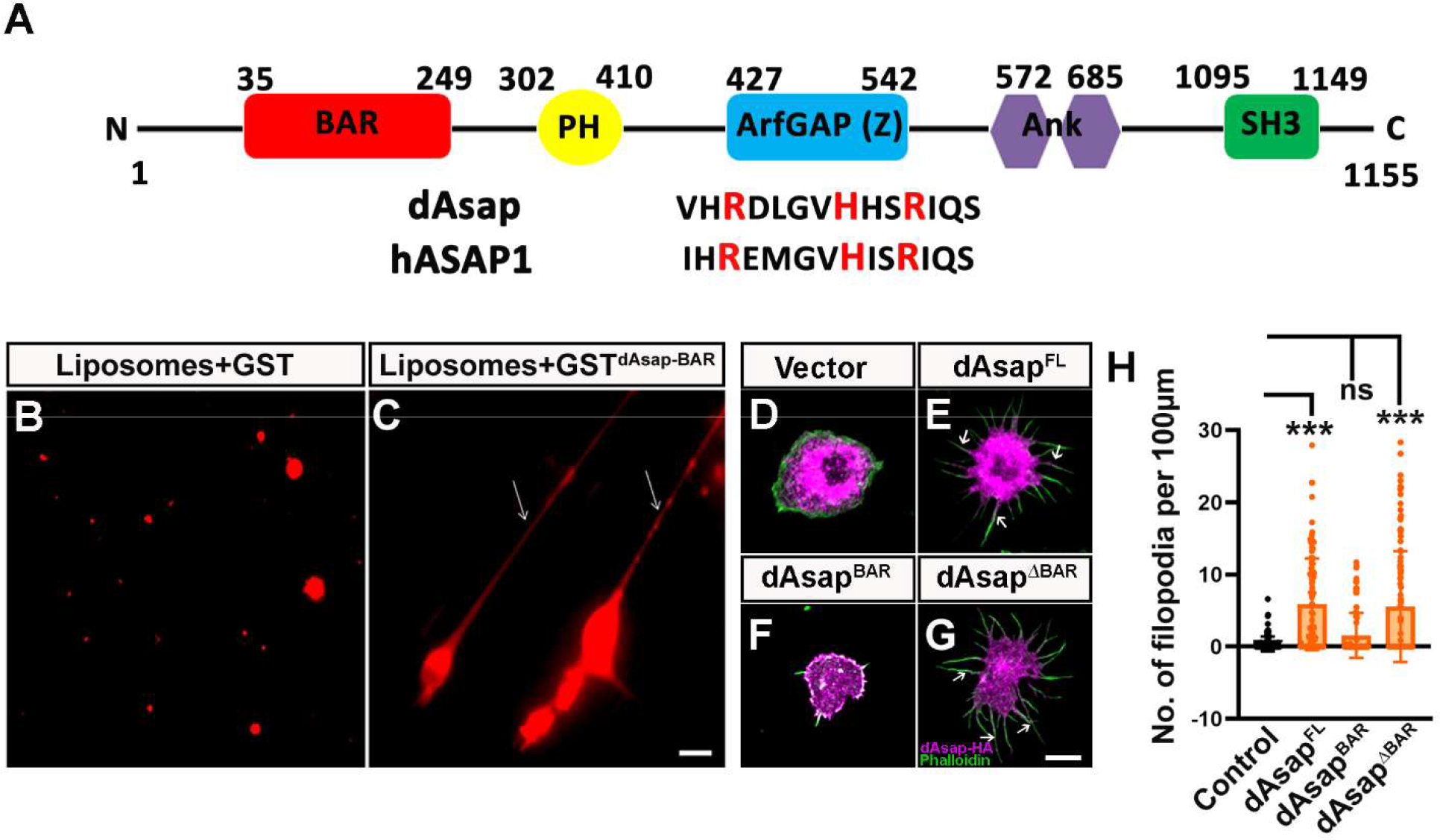
BAR domain of dAsap deforms synthetic liposomes *in vitro* but does not induce membrane tubules in cultured cells. **(A)** Schematic representation of dAsap showing that it contains various conserved domains that can regulate membrane dynamics and Arf-dependent activities. The amino acid residues of the ArfGAP domain (labeled in red) are conserved in humans and *Drosophila*. **(B-C)** The dAsap^BAR^ domain induces tubules when incubated with liposomes. Arrow in C shows tubules formed after 10 minutes of incubation of liposomes with the dAsap^BAR^ domain. Scale bar in C represent 20 µm. **(D-G)** Representative confocal image of S2R+ cells transfected with (D) vector alone, (E) *HA-dAsap^FL^*(aa 1-1155), (F) *HA-dAsap^BAR^*(aa 1-249), and (G) *HA-dAsap^ΔBAR^*(aa 250-1155) co-labeled with anti-HA antibodies (shown in magenta) and FITC-Phalloidin stain (shown in green). Arrows in E and G indicate filopodia. The scale bar in G represents 10 µm. **(H)** Histogram showing quantification of the average number of filopodia per 100 µm circumference in vector alone (0.38 ± 0.10), *HA-dAsap^FL^*(5.86 ± 0.60), *HA-dAsap^BAR^*(1.56 ± 0.30) and *HA-dAsap****^Δ^****^BAR^* (5.56 ± 0.71). \*\*\**p* <0.0001; ns, not significant. The error bar represents the standard error of the mean (SEM); the statistical analysis was done using one-way ANOVA followed by post-hoc Tukey’s multiple-comparison test.

### dAsap functions in neurons to regulate NMJ morphogenesis

Earlier reports have indicated the roles of BAR family proteins in regulating synaptic morphogenesis and function (Elias and Feingold, 1992; Mallik *et al*., 2022; Nahm et al., 2010; Rikhy et al., 2002; Ukken *et al*., 2016; Yong et al., 2020). Our small-scale targeted RNAi-mediated genetic screen in *Drosophila* on BAR domain-containing proteins identified dAsap as a key regulator of synapse morphology (Mallik *et al*., 2017). In order to decipher the molecular mechanism by which dAsap regulates NMJ morphogenesis, we created loss-of-function mutants of *dAsap* using CRISPR/Cas9-mediated genome editing technology. We obtained two viable null mutants, *dAsap^K23^* (4190 bp deletion) and *dAsap^B52^*(4275 bp deletion) (Figure 2A). Semiquantitative RT-PCR and Western blot analysis confirmed that the d*Asap* mutants produced no detectable *dAsap* transcript and were protein null (Figure 2G and Figure S1A-B). To exclude any consequence of background mutation in homozygous mutants, we performed all further experiments in the heteroallelic combination of *dAsap^B52^* and *dAsap^K23^* (*dAsap^K23/B52^*).

**Figure 2.**
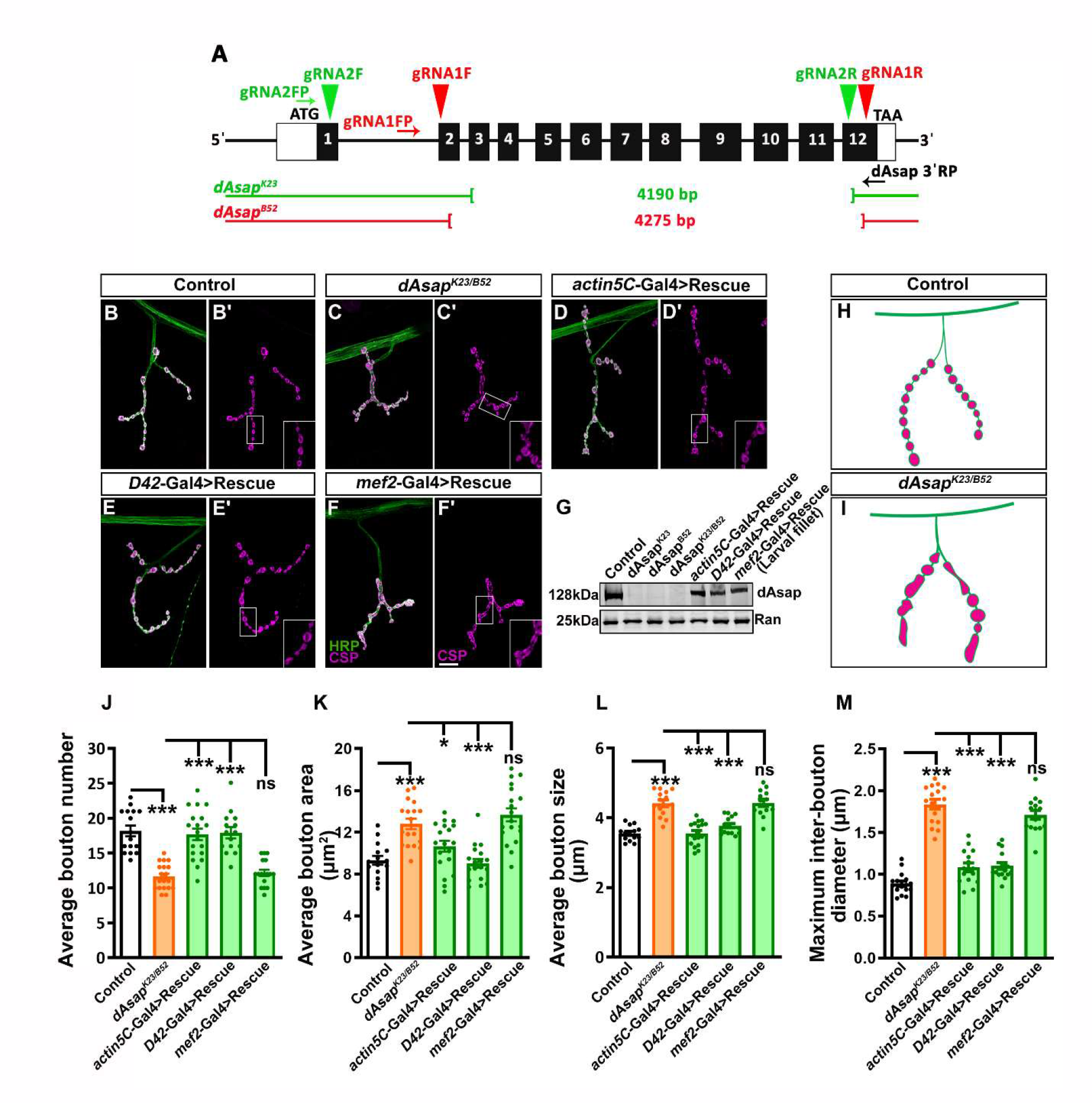
dAsap functions in neurons to regulate NMJ morphogenesis in *Drosophila*. **(A)** Schematic representation of the *dAsap* locus showing exons (solid black boxes, 1-12) and introns (thin black lines). Red and green arrowheads represent gRNA target sites to generate *dAsap* alleles. The first gRNA (red arrowhead) and second gRNA (green arrowhead) were designed for the 2^nd^ and 12^th^ and 1^st^ and 12^th^ exon of the *dAsap* locus, respectively. Mutant lines were screened using Asap 5′ FP and Asap 3′ RP primers (black arrows). The thin green and red line represents *dAsap^K23^* (4190 bp) and *dAsap^B52^* (4275 bp) deletion alleles. **(B-F**′) Confocal images of NMJ synapses at muscle 4 of A2 hemisegment showing synaptic growth in (B-B′) *W^1118^* (Control), (C-C′) *dAsap^B52/K23^* heteroallelic mutant, (D-D′) *actin5C-Gal4* driven rescue line (*actin5C-Gal4, dAsap^K23/B52^; UAS-dAsap^FL^/+*), (E-E′) *D42*-Gal4 driven rescue line (*dAsap^B52/K23^; D42-Gal4/UAS-dAsap^FL^*) and (F-F′) *mef2*-Gal4 driven rescue line (*dAsap^B52/K23^; mef2-Gal4/UAS-dAsap^FL^*) double immunolabeled with neuronal membrane marker, HRP (green) and presynaptic vesicle marker, CSP (magenta). Note that the mutant showed altered NMJ morphology, and these defects were rescued by expressing *UAS-dAsap^FL^* in motor neurons but not in muscles. The scale bar in F′ for (B-F′) represents 20 µm. **(G)** Western blot showing protein levels of dAsap in controls, homozygous *dAsap^K23^*, homozygous *dAsap^B52^*, heteroallelic *dAsap^K23/B52^*, *actin5C*-Gal4 driven rescue, *D42*-Gal4 driven rescue, and *mef2* Gal4 driven rescue animals. The dAsap protein levels were not detected in homozygous and heteroallelic null mutants. Ran protein levels were used as an internal loading control. **(H-I)** Schematic representation of NMJ in control (G) and mutant (H) animals as indicated. Note that the mutants show fused and larger size boutons compared to control NMJ. **(J)** Histogram showing an average number of boutons from muscle 4 NMJ at A2 hemisegment in *W^1118^* control animals (18.19 ± 0.75), *dAsap^K23/B52^* (11.63 ± 0.42), *actin5C-Gal4, dAsap^K23/B52^; UAS-dAsap^FL^/+* (17.68 ± 0.81), *dAsap^B52/K23^; D42-Gal4/UAS-dAsap^FL^* (17.93 ± 0.81), *dAsap^B52/K23^; mef2-Gal4/UAS-dAsap^FL^* (12.18 ± 0.45). *p<0.05, ***p<0.001; ns, not significant. The error bar represents the standard error of the mean (SEM); the statistical analysis was done using one-way ANOVA followed by post-hoc Tukey’s multiple-comparison test. **(K)** Histogram showing an average bouton area from muscle 4 NMJ at A2 hemisegment in control animals (9.30 ± 0.42 µm^2^), *dAsap^K23/B52^* (12.80 ± 0.49 µm^2^), *actin5C-Gal4, dAsap^K23/B52^; UAS-dAsap^FL^/+* (10.66 ± 0.52 µm^2^), *dAsap^B52/K23^; D42-Gal4/UAS-dAsap^FL^* (9.02 ± 0.40 µm^2^), *dAsap^B52/K23^; mef2-Gal4/UAS-dAsap^FL^* (13.66 ± 0.61 µm^2^). *p<0.05, ***p<0.001; ns, not significant. The error bar represents the standard error of the mean (SEM); the statistical analysis was done using one-way ANOVA followed by post-hoc Tukey’s multiple-comparison test. **(L)** Histogram showing an average bouton size from muscle 4 NMJ at A2 hemisegment in *W^1118^* control animals (3.55 ± 0.05 µm), *dAsap^K23/B52^*(4.41 ± 0.09 µm), *actin5C-Gal4, dAsap^K23/B52^; UAS-dAsap^FL^/+* (3.55 ± 0.08 µm), *dAsap^B52/K23^; D42-Gal4/UAS-dAsap^FL^* (3.77 ± 0.06 µm), *dAsap^B52/K23^; mef2-Gal4/UAS-dAsap^FL^* (4.42 ± 0.10 µm). *p<0.05, ***p<0.001; ns, not significant. The error bar represents the standard error of the mean (SEM); the statistical analysis was done using one-way ANOVA followed by post-hoc Tukey’s multiple-comparison test. **(M)** Histogram showing maximum inter-bouton diameter from muscle 4 NMJ at A2 hemisegment in control animals (0.88 ± 0.03 µm), *dAsap^K23/B52^* (1.83 ± 0.05 µm), *actin5C-Gal4, dAsap^K23/B52^; UAS-dAsap^FL^/+* (1.08 ± 0.05 µm), *dAsap^B52/K23^; D42-Gal4/UAS-dAsap^FL^* (1.10 ± 0.03 µm), *dAsap^B52/K23^; mef2-Gal4/UAS-dAsap^FL^* (1.71 ± 0.05 µm). *p<0.05, ***p<0.001; ns, not significant. The error bar represents the standard error of the mean (SEM); the statistical analysis was done using one-way ANOVA followed by post-hoc Tukey’s multiple-comparison test.

To quantify the NMJ growth phenotypes in *dAsap* mutants, we measured the average bouton number, bouton area, bouton size, and maximum inter-bouton diameter in the third instar larval NMJ synapse at muscle 4 of the A2 hemisegment. We found that, compared to the control synapse, the *dAsap* mutant showed a significant reduction in average bouton number (control, 18.19 ± 0.75; *dAsap^K23/B52^*, 11.63 ± 0.42, *p*<0.0001), increased in average bouton area (control, 9.30 ± 0.42 µm^2^; *dAsap^K23/B52^*, 12.80 ± 0.49 µm^2^, *p*<0.0001), average bouton size (control, 3.55 ± 0.05 µm; *dAsap^K23/B52^*, 4.41 ± 0.09 µm, *p*<0.0001) and maximum inter-bouton diameter (control, 0.88 ± 0.03 µm; *dAsap^K23/B52^*, 1.83 ± 0.05 µm, *p*<0.0001) (Figure 2B-M and Figure S1).

The aberrant synaptic morphology of *dAsap* mutants was fully restored by expressing a full-length *dAsap* transgene in the motor neurons (*dAsap^K23/B52^; UAS-dAsap^FL^/D42-Gal4*) or ubiquitously (a*ctin5C-Gal4, dAsap^K23/B52^; UAS-dAsap^FL^/+*) in *dAsap^K23/B52^* mutants. However, muscle-specific expression of a full-length *dAsap* transgene in *dAsap^K23/B52^* mutant (*dAsap^K23/B52^; UAS-dAsap^FL^/mef2-Gal4*) did not restore the NMJ defects (Figure 2B-M). These data suggest that dAsap is required in neurons to regulate synapse morphology in *Drosophila*.

### dAsap regulates microtubule organization but not actin-based cytoskeleton at the synapse

The microtubule-based cytoskeleton is crucial for regulating various aspects of neuronal morphogenesis, including axonal growth, growth cone guidance, stabilization of dendrites, and formation of the nascent synapse (Choudhury et al., 2016; Roos et al., 2000; Wechsler et al., 2018). Several studies have implicated the role of ASAP1 as an essential component in modulating cytoskeletal dynamics, including the organization of actin filaments and stress fiber formation in the cells (Chen *et al*., 2020; Gasilina *et al*., 2019). To assess whether defects in synapse morphology in *dAsap* mutants are associated with the disorganized cytoskeleton at the NMJs, we immunostained *dAsap^K23/B52^*NMJ for microtubule-associated protein 1B (MAP1B/Futsch) using 22C10 antibody. Interestingly, we observed a significant reduction in the percentage of Futsch-positive loops in *dAsap* mutants compared to the control NMJs (control, 35.49 ± 2.40 %; *dAsap^K23/B52^*, 23.61 ± 2.40 %, *p*=0.002) (Figure 3A-D). The reduced microtubule loops in mutants were fully rescued to the control level by expressing a full-length *dAsap* transgene in the motor neurons (*dAsap^K23/B52^; UAS-dAsap^FL^/D42-Gal4*, 39.21 ± 2.68 %, *p*=0.0003) of *dAsap^K23/B52^*mutants (Figure 3A-D). In contrast, *dAsap* mutants did not show an altered number of moesin-GFP punctae, suggesting that actin-based cytoskeleton was not impaired in *dAsap* mutants at the NMJ (control, 0.18 ± 0.01; *dAsap^K23/B52^*; *UAS-moesinGFP/D42-Gal4*, 0.19 ± 0.02, *p*=0.769) (Fig 3E-H). These data indicate that dAsap-dependent microtubule loop organization may be crucial for NMJ organization.

**Figure 3.**
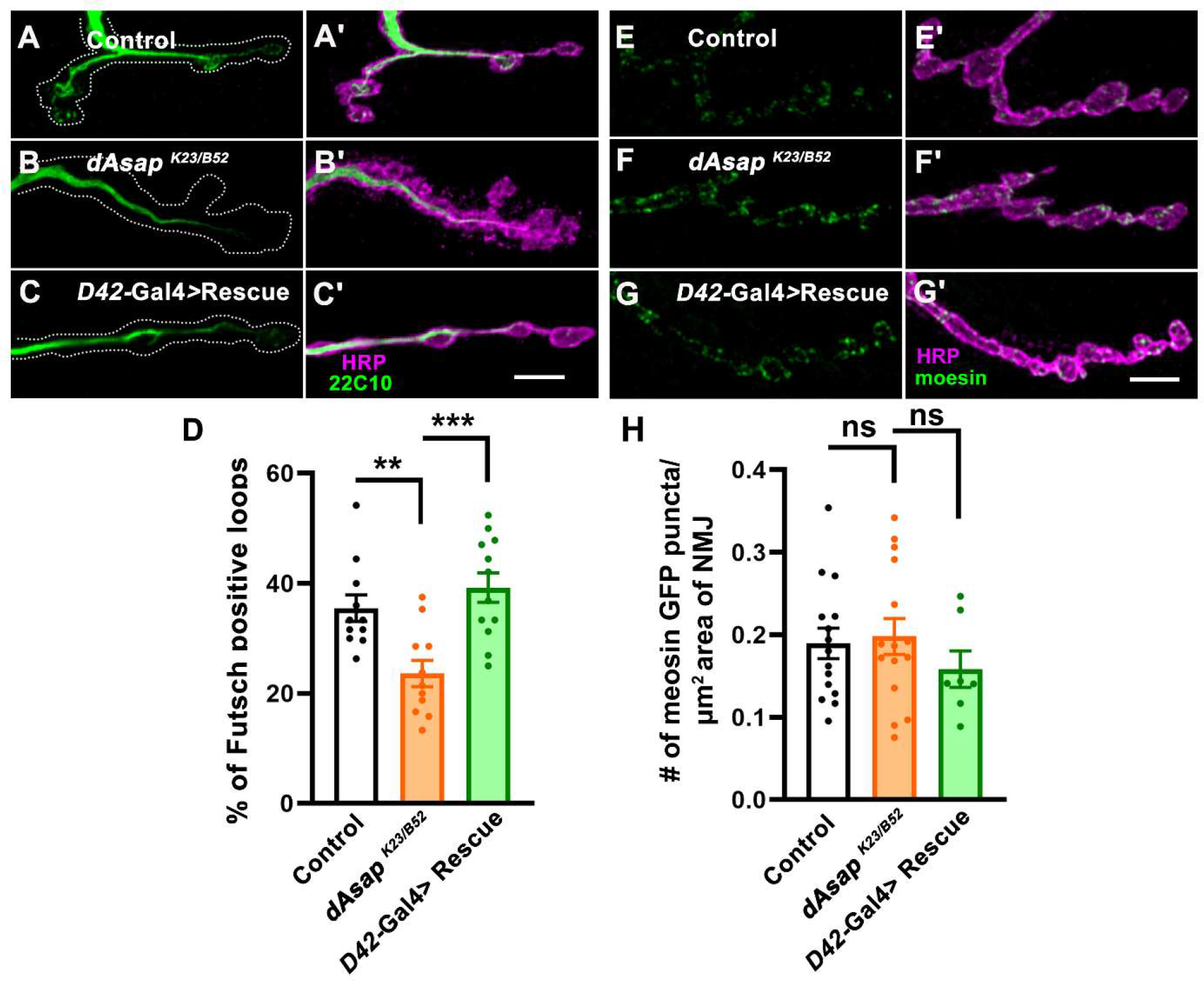
*dAsap* mutation affects microtubule-based cytoskeleton at NMJ. **(A-C**′) Confocal images of NMJ synapses at muscle 4 of A2 hemisegment showing futsch loops in (A, A′) control, (B, B′) *dAsap^B52/K23^* heteroallelic mutant and (C, C′) *D42*-Gal4 driven rescue line (*dAsap^B52/K23^; D42-Gal4/UAS-dAsap^FL^*) double immunolabeled with 22C10 (green) and a neuronal membrane marker, HRP (magenta). Futsch positive loops were reduced in ASAP mutant. Scale bar in C′ (for A-C′) represent 4 μm. **(D)** Histogram showing the percentage of the futsch positive loops from muscle 4 NMJ at A2 hemisegment in control animals (35.49 ± 2.409 %), *dAsap^K23/B52^* (23.61 ± 2.403 %) and *dAsap^B52/K23^; D42-Gal4/UAS-dAsap^FL^* (39.21 ± 2.686 %). **p<0.01, ***p<0.001; ns, not significant. The error bar represents the standard error of the mean (SEM); the statistical analysis was done using one-way ANOVA followed by post-hoc Tukey’s multiple-comparison test. **(E-G**′) Confocal images of NMJ synapses at muscle 4 of A2 hemisegment showing Moesin-GFP punctae in (E, E′) *D42-Gal4/UAS-moesinGFP* (control), (F, F′) *dAsap^K23/B52^; D42-Gal4/UAS-moesinGFP* and (G, G′) *dAsap^K23/B52^; D42-Gal4, UAS-moesinGFP/D42-Gal4, UAS-dAsap^FL^* double immunolabeled with anti-GFP (green) and HRP (magenta). Scale bar in G′ (for E-G′) represent 20 μm. **(H)** Histogram showing the number of moesin-GFP punctae per area of NMJ from muscle 4 at A2 hemisegment in *D42-Gal4/UAS-moesinGFP* (control) animals (0.18 ± 0.01), *dAsap^K23/B52^; D42-Gal4/UAS-moesinGFP* (0.19 ± 0.02) and *dAsap^K23/B52^; D42-Gal4, UAS-moesinGFP/D42-Gal4, UAS-dAsap^FL^* (0.15 ± 0.02). ns, not significant. The error bar represents the standard error of the mean (SEM); the statistical analysis was done using one-way ANOVA followed by post-hoc Tukey’s multiple-comparison test.

### *dAsap* mutants show elevated miniature frequency and evoked activity at the NMJ

At the neuromuscular junction, Futsch affects synaptic morphology and glutamate release to maintain normal synaptic activity (Lepicard et al., 2014; Roos *et al*., 2000). Consistent with these prior observations, we found that loss of *dAsap* in neurons leads to impaired synapse growth and organization of Futsch-positive loops at the NMJ (Figure 2 and Figure 3). To assess whether impaired synapse organization in *dAsap* mutants is also associated with altered synaptic transmission, we performed electrophysiological measurements in control and *dAsap^K23/B52^* mutant animals. While the amplitude of spontaneous miniature postsynaptic potentials (mEPSP) was not altered, the miniature frequency and excitatory postsynaptic potentials (EPSP) were significantly increased in the mutants compared to control animals (mEPSP amplitude: control, 0.60 ± 0.04 mV; *dAsap^K23/B52^*, 0.73 ± 0.04 mV, *p*=0.066), (mEPSP frequency: control, 1.16 ± 0.17 Hz; *dAsap^K23/B52^*, 4.72 ± 0.56 Hz, *p*<0.0001) and (EPSP amplitude: control, 51.45 ± 1.27 mV; *dAsap^K23/B52^*, 60.10 ± 2.48 mV, *p*=0.004), (Figure 4A-I). However, *dAsap* mutants showed no significant change in the quantal content compared to control animals (QC: control, 88.60 ± 5.27; *dAsap^K23/B52^*, 83.64 ± 3.97, *p*=0.4681) (Figure 4J).

**Figure 4.**
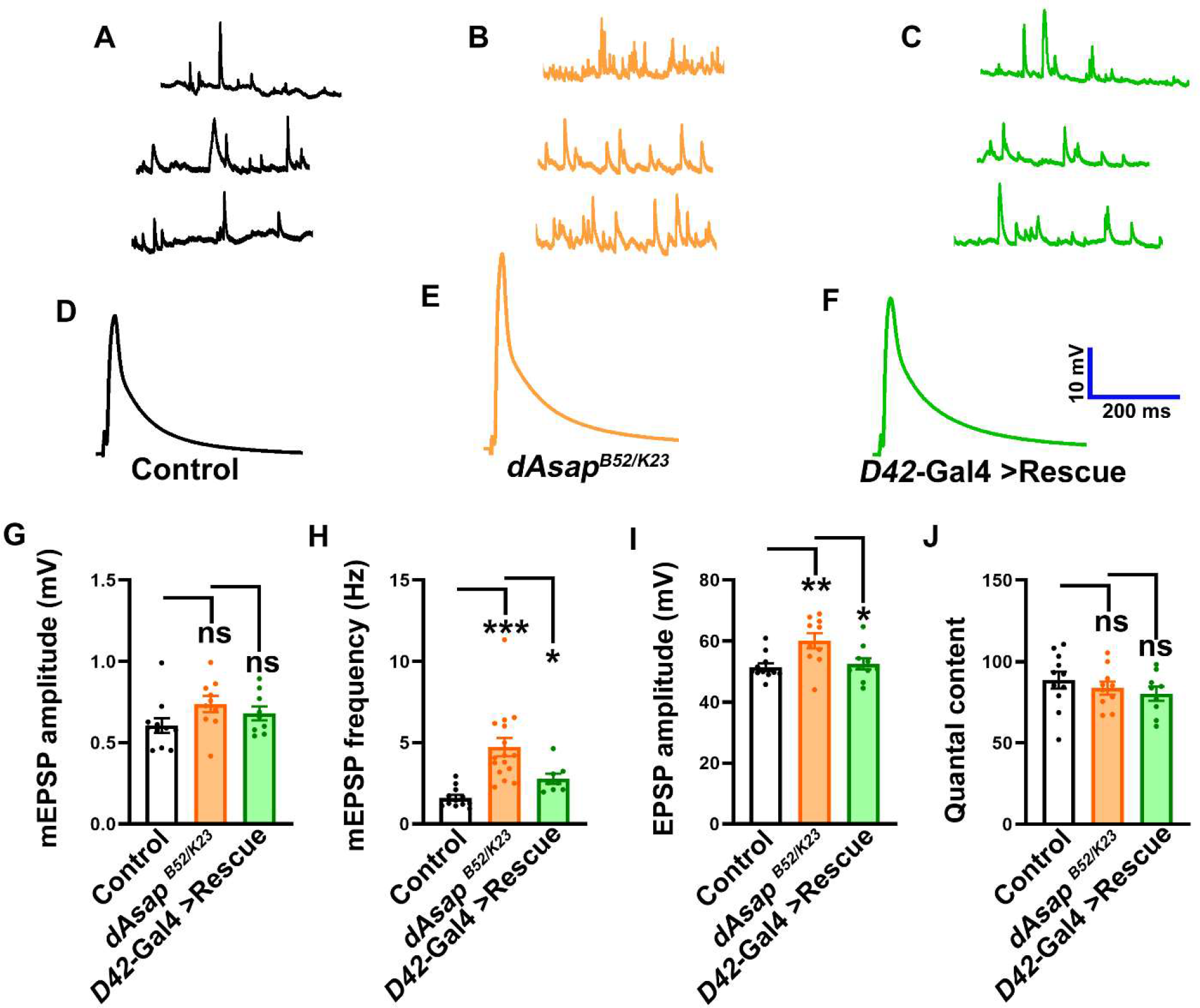
*dAsap* mutants show increased mEPSP frequency and EPSP amplitude. **(A-C)** Representative traces of mEPSP in (A) control, (B) *dAsap^K23/B52^*, and (C) *dAsap^B52/K23^; D42-Gal4/UAS-dAsap^FL^* larvae. **(D-F)** Representative traces of EPSP in (D) control, (E) *dAsap^K23/B52^*, (F) *dAsap^B52/K23^; D42-Gal4/UAS-dAsap^FL^* larvae. **(G)** Histogram showing mEPSP amplitude from muscle 6/7 NMJ at A2 hemisegment in control animals (0.60 ± 0.04 mV), *dAsap^K23/B52^*(0.73 ± 0.04 mV), *dAsap^B52/K23^; D42-Gal4/UAS-dAsap^FL^* (0.68 ± 0.04 mV). ns, not significant. The error bar represents the standard error of the mean (SEM); the statistical analysis was done using one-way ANOVA followed by post-hoc Tukey’s multiple-comparison test. **(H)** Histogram showing mEPSP frequency from muscle 6/7 NMJ at A2 hemisegment in control animals (1.16 ± 0.17 Hz), *dAsap^K23/B52^* (4.72 ± 0.56 Hz), *dAsap^B52/K23^; D42-Gal4/UAS-dAsap^FL^* (2.77 ± 0.31 Hz). *p<0.05, ***p<0.001. The error bar represents the standard error of the mean (SEM); the statistical analysis was done using one-way ANOVA followed by post-hoc Tukey’s multiple-comparison test. **(I)** Histogram showing EPSP amplitude from muscle 6/7 NMJ at A2 hemisegment in control animals (51.45 ± 1.27 mV), *dAsap^K23/B52^* (60.10 ± 2.48 mV), *dAsap^B52/K23^; D42-Gal4/UAS-dAsap^FL^* (52.47 ± 1.82 mV). *p<0.05, **p<0.01. The error bar represents the standard error of the mean (SEM); the statistical analysis was done using one-way ANOVA followed by post-hoc Tukey’s multiple-comparison test. **(J)** Histogram showing quantification of quantal content in control animals (88.60 ± 5.27), *dAsap^K23/B52^* (83.64 ± 3.97), *dAsap^B52/K23^; D42-Gal4/UAS-dAsap^FL^* (80.27 ± 4.36). ns, not significant. The error bar represents the standard error of the mean (SEM); the statistical analysis was done using one-way ANOVA followed by post-hoc Tukey’s multiple-comparison test.

The increased miniature frequency and evoked amplitude in *dAsap* mutants were significantly rescued by expressing a full-length *dAsap* transgene in the motor neurons (mEPSP frequency: *dAsap^K23/B52^*, 4.72 ± 0.56 Hz; *dAsap^K23/B52^; UAS-dAsap^FL^/D42-Gal4*: 2.77 ± 0.31 Hz, *p*=0.028; EPSP amplitude: *dAsap^K23/B52^*, 60.10 ± 2.48 mV, *dAsap^K23/B52^; UAS-dAsap^FL^/D42-Gal4,* 52.47 ± 1.82 mV, *p*=0.023; (Figure 4). Collectively, these data indicate that loss of dAsap in neurons results in gains of neurotransmission at the *Drosophila* NMJs.

### Loss of *dAsap* increases synaptic Brp punctae and synaptic vesicle release

Our electrophysiological analysis of *dAsap* mutants revealed a higher miniature frequency than the control NMJs. These physiological changes in mutants could arise due to altered active zone numbers or changes in the organization of glutamate receptors at synapses. To address this, we labeled the NMJs with anti-Bruchpilot (Brp) and anti-GluRIII antibodies to quantify the number of the presynaptic active zone and postsynaptic glutamate receptors. We found a significant increase in the number of Brp punctae per NMJ in *dAsap* mutants compared to the control NMJs (average number of Brp punctae/NMJ: control, 404.2 ± 19.09; *dAsap^K23/B52^*, 625.9 ± 39.98, *p*=0.0003). Moreover, we found increased Brp punctae density at the mutant NMJ when compared to controls (average BRP density: control, 1.79 ± 0.05 µm^2^; *dAsap^K23/B52^*, 2.20 ± 0.06 µm^2^, *p*<0.0001) (Figure 5A-L). Additionally, the area of the GluR cluster was increased in *dAsap* mutants compared to control animals (control, 0.48 ± 0.03 µm^2^; *dAsap^K23/B52^*, 1.29 ± 0.06 µm^2^, *p*<0.0001) (Figure 5A-L). Moreover, the gross organization of GluRIII clusters appears altered and tightly apposed to the corresponding Brp-positive active zone (Figure 5A-I). The increased Brp density and GluRIII receptor clusters in mutants were fully restored to the wild-type levels by expressing a full-length *dAsap* transgene in the motor neurons of *dAsap* mutants (average BRP density: *dAsap^K23/B52^*, 2.20 ± 0.06 µm^2^; *dAsap^K23/B52^; UAS-dAsap^FL^/D42-Gal4*, 1.93 ± 0.06 µm^2^, *p*=0.005; the average number of BRP puncta/NMJ: *dAsap^K23/B52^*, 625.9 ± 39.98; *dAsap^K23/B52^; UAS-dAsap^FL^/D42-Gal*, 447.6 ± 14.90, *p*=0.002) (Figure 5J-L).

**Figure 5.**
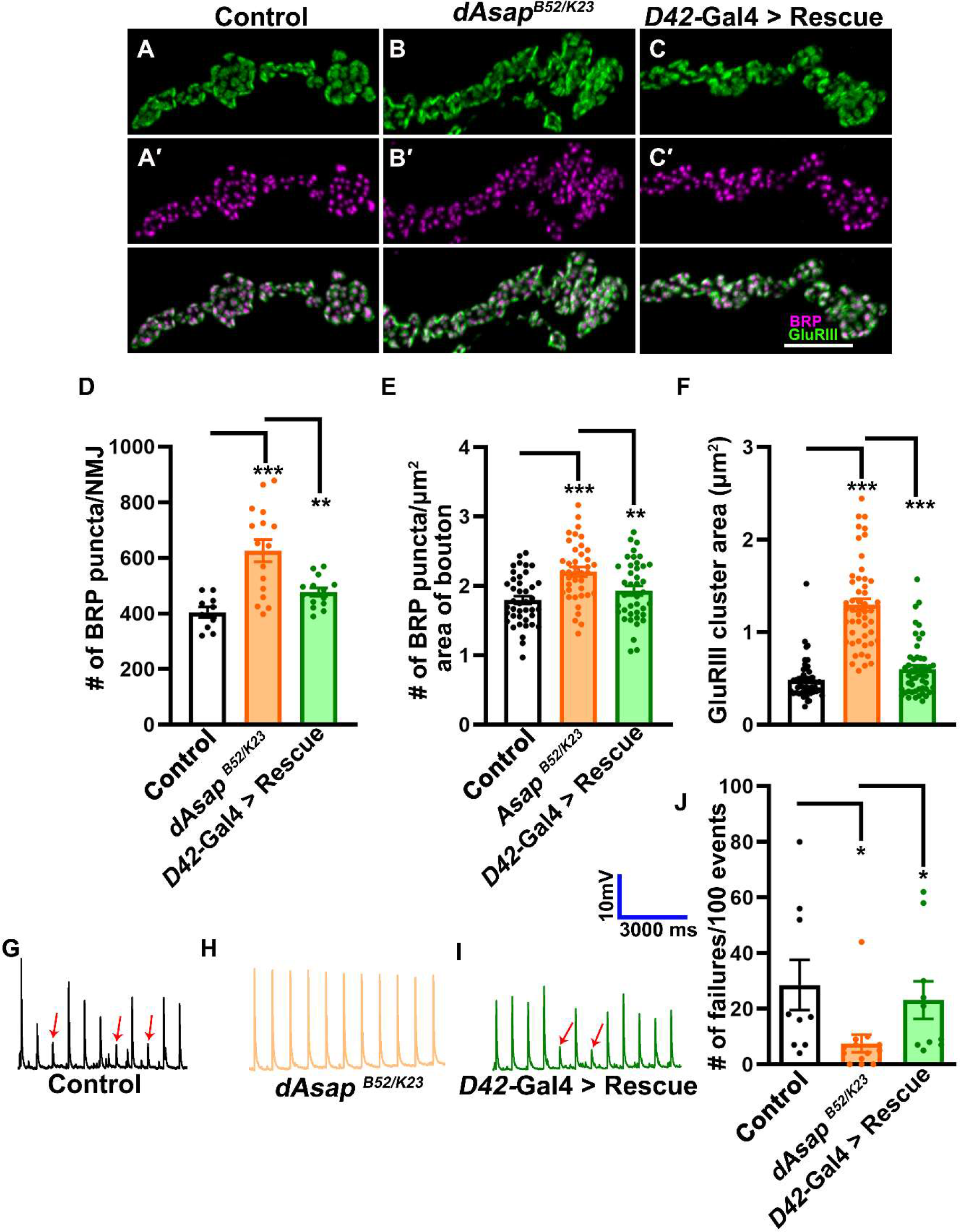
*dAsap* mutation shows more active zone numbers at the NMJs. **(A-I)** Confocal images of NMJ synapses at muscle 4 of A2 hemisegment of control, *dAsap^K23/B52^*mutants and (C) *dAsap^B52/K23^; D42-Gal4/UAS-dAsap^FL^* double immunolabeled with antibodies against active zones marker Bruchpilot, Brp (A-C, magenta) and GluRIII (D-F, green). Note that *dAsap* mutation shows an increased active zone number. The scale bar in I represents 2.5 μm. **(J)** Histogram showing the number of Brp punctae per NMJ from muscle 4 NMJ at A2 hemisegment in control animals (404.2 ± 19.09), *dAsap^K23/B52^* (625.9 ± 39.98), *dAsap^B52/K23^; D42-Gal4/UAS-dAsap^FL^* (476.6 ± 14.90). **p<0.01, ***p<0.001. The error bar represents the standard error of the mean (SEM); the statistical analysis was done using one-way ANOVA followed by post-hoc Tukey’s multiple-comparison test. **(K)** Histogram showing the number of Brp punctae per μm^2^ area of bouton from muscle 4 NMJ at A2 hemisegment in control animals (1.79 ± 0.05 μm^2^), *dAsap^K23/B52^* (2.20 ± 0.06 μm^2^), *dAsap^B52/K23^; D42-Gal4/UAS-dAsap^FL^* (1.93 ± 0.06 μm^2^). **p<0.01, ***p<0.001. The error bar represents the standard error of the mean (SEM); the statistical analysis was done using one-way ANOVA followed by post-hoc Tukey’s multiple-comparison test. **(L)** Histogram showing the area of GluRIII cluster from muscle 4 NMJ at A2 hemisegment in control animals (0.48 ± 0.03 μm^2^), *dAsap^K23/B52^* (1.29 ± 0.06 μm^2^), *dAsap^B52/K23^; D42-Gal4/UAS-dAsap^FL^* (0.60 ± 0.04 μm^2^). ***p<0.001. The error bar represents the standard error of the mean (SEM); the statistical analysis was done using one-way ANOVA followed by post-hoc Tukey’s multiple-comparison test. **(M-O)** Representative traces of action potential firing at 0.1 mM extracellular Ca^2+^ in (M) control, (N) *dAsap^K23/B52^*, (O) *dAsap^B52/K23^; D42-Gal4/UAS-dAsap^FL^* larvae. **(P)** Histogram showing percentage of failures in control animals (28.56 ± 9.04 %), *dAsap^K23/B52^* (7.53 ± 3.17 %), *dAsap^B52/K23^; D42-Gal4/UAS-ASAP^FL^* (23.10 ± 6.71 %). *p<0.05. The error bar represents the standard error of the mean (SEM); the statistical analysis was done using one-way ANOVA followed by post-hoc Tukey’s multiple-comparison test.

Since we found an increased number of Brp punctae in the *dAsap* mutants, we hypothesized that the mutant synapse would be able to sustain synaptic vesicle release under low extracellular calcium. To test this hypothesis, we used failure analysis to determine the number of failed release events and assess presynaptic release probability at low (0.1 mM) extracellular Ca^2+^ concentrations. Interestingly, we observed significantly low failure probability in *dAsap* mutants when compared to control NMJs (control, 28.56 ± 9.04 %; *dAsap^K23/B52^*, 7.53 ± 3.17 %, *p*=0.023) (Figure 5M-P). Moreover, expressing a full-length *dAsap* transgene in *dAsap^K23/B52^* mutants significantly restored the synaptic failures (*dAsap^K23/B52^*, 7.53 ± 3.17 %; *dAsap^K23/B52^; UAS-dAsap^FL^/D42-Gal4*: 23.10 ± 6.71 %, *p*=0.034) (Figure 5M-P). Taken together, these data suggest that *dAsap* mutants have a high release probability, possibly due to an increased number of synaptic vesicle release sites.

### dAsap regulates calcium release at synapses to maintain robust neurotransmission

The structural and biochemical studies revealed the functional requirement of calcium in the regulation of ASAP-mediated GTP hydrolysis, further suggesting a link between ASAP and Arf6-mediated calcium signaling (Ismail *et al*., 2010). Besides, Arf6-GTP has been implicated in phospholipase C-inositol-3-phosphate (PLC-IP3) mediated calcium release during acrosomal exocytosis (Pelletan *et al*., 2015). Similarly, in neurons, vesicle fusion is highly orchestrated and involves calcium influx, preparation of fusion machinery, and exocytosis of synaptic vesicles. Hence, we speculated that enhanced evoked responses in *dAsap* mutant could arise due to: a) higher active zone number and/or b) the release of more calcium from intracellular stores, promoting more vesicle fusion at the synaptic terminals. Consistent with this hypothesis, our analysis in *dAsap* mutants revealed an increased active zone number, promoting more vesicle fusion (Figure 5).

Next, we assessed calcium release from intracellular stores in *dAsap* mutants. We found that pharmacological or genetic perturbation of IP3R (blocked using neuronally expressed *ip3-sponge.m30*) and RyR (blocked using Dantrolene) did not show significant changes in evoked activity compared to the control (Figure 6E, L-M). Interestingly, blocking IP3R and RyR in *dAsap* mutants suppressed the enhanced evoked activity to the control levels (control, 39.24 ± 1.33 mV; *D42-Gal4/UAS-ip3-sponge.m30*+Dant, 40.39 ± 0.90 mV, *p*=0.485; *dAsap^K23/B52^*, 45.17 ± 1.11 mV, *p*=0.003; *dAsap^K23/B52^*; *D42-Gal4/UAS-ip3-sponge.m30* + Dantrolene, 37.44 ± 1.22 mV, *p*=0.0002) (Figure 6). However, miniature frequencies were not restored following pharmacological or genetic blockade of RyR in *dAsap* mutants (control, 2.11 ± 0.21 Hz; *D42-Gal4/UAS-ip3-sponge.m30*+Dant, 1.09 ± 0.11 Hz, *p*=0.0005; *dAsap^K23/B52^*, 2.91 ± 0.27 Hz, *p*=0.036; *dAsap^K23/B52^*; *D42-Gal4/UAS-ip3-sponge.m30* + Dantrolene, 2.67 ± 0.57 Hz, *p*=0.706) (Figure 6).

**Figure 6.**
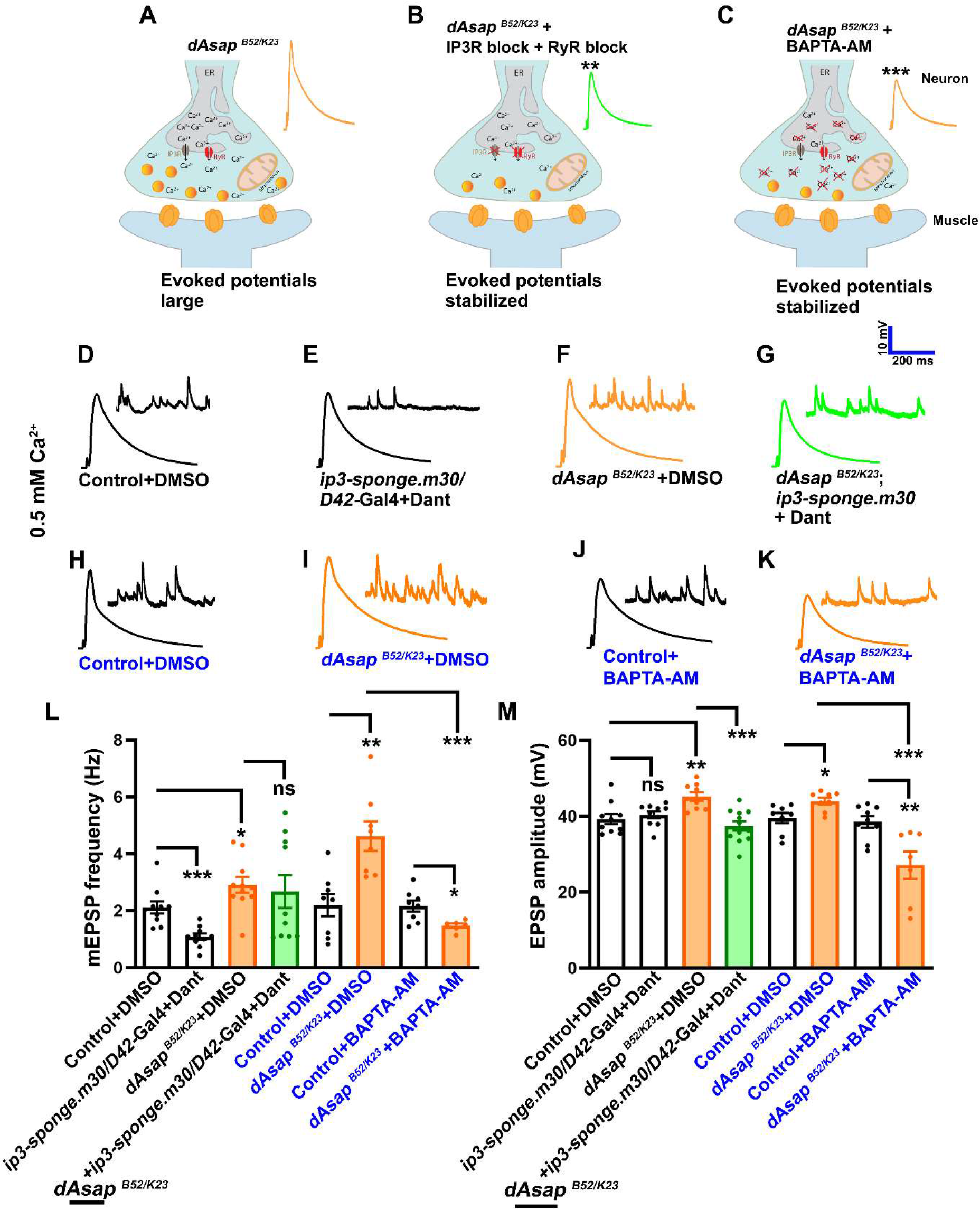
dAsap regulates synaptic calcium release from intracellular stores. **(A-C)** Schematic representation of the role of IP3 receptor (IP3R), Ryanodine receptor (RyR) in the endoplasmic reticulum, mitochondrial calcium uniporter (MCU), Ca_v_2 calcium channel, and synaptic vesicles at the presynaptic terminals. **(D-G)** Representative traces of mEPSPs and EPSPs at 0.5 mM Ca^2+^ in (D) control + DMSO, (E) *D42-Gal4/UAS-ip3-sponge.m30*+Dant, (F) *dAsap^K23/B52^* + DMSO, (G) *dAsap^K23/B52^; D42-Gal4/UAS-ip3-sponge.m30* + Dantrolene larvae. **(H-K)** Representative traces of mEPSPs and EPSPs at 0.5 mM Ca^2+^ in (H) control + DMSO, (I) *dAsap^K23/B52^* + DMSO, (J) control + BAPTA-AM, (K) *dAsap^K23/B52^* + BAPTA-AM larvae. **(L)** Histogram showing mEPSP frequency from muscle 6/7 NMJ at A2 hemisegment in control + DMSO (2.11 ± 0.21 Hz), *D42-Gal4/UAS-ip3-sponge.m30*+Dant (1.09 ± 0.11 Hz), *dAsap^K23/B52^* + DMSO (2.91 ± 0.27 Hz), *dAsap^K23/B52^; D42-Gal4/UAS-ip3-sponge.m30* + Dantrolene (2.67 ± 0.57 Hz), control + DMSO (2.20 ± 0.39 Hz), *dAsap^K23/B52^* + DMSO (4.61 ± 0.51 Hz), control + BAPTA-AM (2.16 ± 0.19 Hz), *dAsap^K23/B52^* + BAPTA-AM (1.47 ± 0.07 Hz). The error bar represents the standard error of the mean (SEM); the statistical analysis was done using one-way ANOVA followed by post-hoc Tukey’s test. *p<0.05, **p<0.01, ***p<0.001; ns, not significant. **(M)** Histogram showing EPSP amplitude from muscle 6/7 NMJ at A2 hemisegment in control + DMSO (39.24 ± 1.33 mV), *D42-Gal4/UAS-ip3-sponge.m30*+Dant (40.39 ± 0.90 mV), *dAsap^K23/B52^* + DMSO (45.17 ± 1.11 mV), *dAsap^K23/B52^; D42-Gal4/UAS-ip3-sponge.m30* + Dantrolene (37.44 ± 1.22 mV), control + DMSO (39.54 ± 1.34 mV), *dAsap^K23/B52^* + DMSO (43.99 ± 0.90 mV), control + BAPTA-AM (38.52 ± 1.48 mV), *dAsap^K23/B52^* + BAPTA-AM (27.13 ± 3.58 mV). The error bar represents the standard error of the mean (SEM); the statistical analysis was done using one-way ANOVA followed by post-hoc Tukey’s test. *p<0.05, **p<0.01, ***p<0.001; ns, not significant.

In order to further test the idea that synaptic calcium is elevated in *dAsap* mutants, we chelated the intracellular calcium using membrane-permeable BAPTA-AM. Consistent with the genetic and pharmacological evidence, we found that chelating calcium intracellularly suppressed enhanced evoked activity and miniature frequencies in *dAsap* mutants compared to the DMSO-treated mutant animals (EPSP: *dAsap^K23/B52^*, 43.99 ± 0.90 mV; *dAsap^K23/B52^*+ BAPTA-AM, 27.13 ± 3.58 mV, *p*=0.006; mEPSP frequency: *dAsap^K23/B52^*, 4.61 ± 0.51 Hz; *dAsap^K23/B52^*+ BAPTA-AM, 1.47 ± 0.07 Hz, *p*=0.013). Together, these results are consistent with the idea that elevated intracellular calcium release from ER and increased active zones in *dAsap* mutants contribute to maintaining robust neurotransmission.

### Expression of a dominant negative Arf6 in *dAsap* mutants restores morphological and functional defects

Mammalian ASAP1 and ASAP2 function as Arf effectors through the conserved Arf GAP domain activity (Kam et al., 2000). First, to test if the Arf-GAP domain of dAsap is biochemically conserved, we performed a malachite green colorimetric assay (Chang et al., 2008). The assay revealed that compared to dArf6-GTP, dArf1-GTP was the preferred substrate for the Arf GAP domain of dAsap for GTP hydrolysis (Figure 7A-B). In order to understand the neuronal relevance of the Arf GAP activities on Arf1 and Arf6, we performed genetic interaction studies of *dAsap* mutants with flies expressing dominant negative Arf1 or Arf6. We found that expression of Arf1^DN^ in the motor neurons (*dAsap^K23/B52^; UAS-Arf1^DN^/D42-Gal4*) or ubiquitously (*actin5C-Gal4, dAsap^K23/B52^; UAS-Arf1^DN^/+*) in *dAsap* mutant did not yield viable third instar larvae at 22°C. Moreover, we found that overexpression of Arf1^DN^ but not Arf6^DN^ in motor neurons induced synaptic growth defects (Supplemental Figure S2). Hence, we could not assess epistatic interactions between Arf1^DN^ and *dAsap* mutants. Interestingly, we found that expressing a dominant negative form of Arf6 either in the motor neurons (*dAsap^K23/B52^*, 13.60 ± 0.63; *dAsap^K23/B52^; UAS-Arf6^DN^/D42-Gal4*, 18.00 ± 1.00, *p*<0.0001) or ubiquitously (a*ctin5C-Gal4, dAsap^K23/B52^; UAS-Arf6^DN^/+*, 20.29 ± 0.77, *p*<0.0001) restored NMJ structural defects in *dAsap* mutants (Figure 7E-E′, F-F′, J-L,). However, expression of *Arf6^DN^* in the muscle (*dAsap^K23/B52^; UAS-Arf6^DN^/mef2-Gal4*, 13.87 ± 0.82, *p*=0.798) did not restore the aberrant synapse morphology in *dAsap* mutant but restored the inter-bouton diameter (Figure 7G-G′, J-L). These epistatic interaction studies suggest that dAsap regulates synapse morphology primarily through the Arf6-dependent signaling pathway in *Drosophila*.

**Figure 7.**
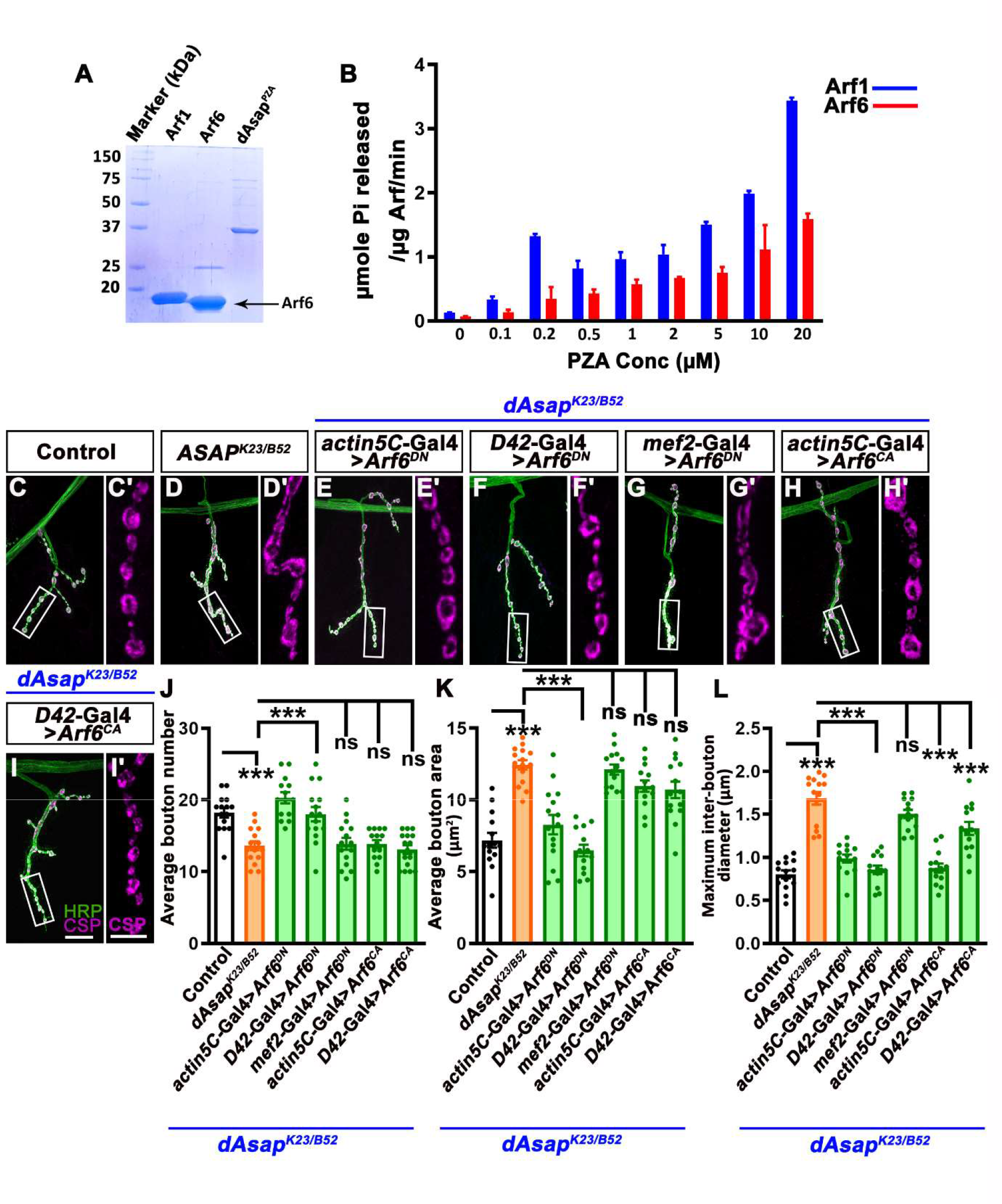
Expression of a Arf6^DN^ rescues developmental defects of *dAsap* mutation. **(A)** Representative gel image stained with Coomassie brilliant blue showing purified proteins, Arf1, Arf6, and dAsap^PZA^ used for the biochemical assay. **(B)** The PZA domain of dAsap has more affinity for Arf1 than Arf6 to hydrolyze the GTP. Arf1 loaded with GTP show more inorganic phosphate release with increasing concentration of PZA. **(C-I**′) Confocal images of NMJ synapses at muscle 4 of A2 hemisegment showing synaptic growths in (C, C′) control, (D, D′) *dAsap^K23/B52^*, (E, E′) *actin5C-Gal4, dAsap^K23/B52^; UAS-Arf6^DN^/+*, (F, F′) *dAsap^K23/B52^; D42-Gal4/UAS-Arf6^DN^*, (G, G′) *dAsap^K23/B52^; mef2-Gal4/UAS-Arf6^DN^*, (H, H′) *actin5C-Gal4, dAsap^K23/B52^; UAS-Arf6^CA^/+* (I, I′) *dAsap^K23/B52^; D42-Gal4/UAS-Arf6^CA^* double immunolabeled with a presynaptic vesicle marker, CSP (magenta) and a neuronal membrane marker, HRP (green). Scale bar in I (for merged images, C-I) and I′ (for C′-I′) represent 10 μm. **(J)** Histogram showing an average number of boutons from muscle 4 NMJ at A2 hemisegment in control animals (18.20 ± 0.66), *dAsap^K23/B52^* (13.60 ± 0.63), *actin5C-Gal4, dAsap^K23/B52^; UAS-Arf6^DN^/+*(20.29 ± 0.77), *dAsap^K23/B52^; D42-Gal4/UAS-Arf6^DN^* (18.00 ± 1.00), *dAsap^K23/B52^; mef2-Gal4/UAS-Arf6^DN^* (13.87 ± 0.82), *actin5C-Gal4, dAsap^K23/B52^; UAS-Arf6^CA^/+* (13.86 ± 0.54), *dAsap^K23/B52^; D42-Gal4/UAS-Arf6^CA^* (13.06 ± 0.58). The error bar represents the standard error of the mean (SEM); the statistical analysis was done using one-way ANOVA followed by post-hoc Tukey’s test. *p<0.05, **p<0.01, ***p<0.001; ns, not significant. **(K)** Histogram showing average bouton area from muscle 4 NMJ at A2 hemisegment in control animals (7.18 ± 0.51µm^2^), *dAsap^K23/B52^* (12.44 ± 0.33 µm^2^), *actin5C-Gal4, dAsap^K23/B52^; UAS-Arf6^DN^/+* (8.27 ± 0.67 µm^2^), *dAsap^K23/B52^; D42-Gal4/UAS-Arf6^DN^* (6.46 ± 0.40 µm^2^), *dAsap^K23/B52^; mef2-Gal4/UAS-Arf6^DN^* (12.12 ± 0.35 µm^2^), *actin5C-Gal4, dAsap^K23/B52^; UAS-Arf6^CA^/+* (10.96 ± 0.39 µm^2^), *dAsap^K23/B52^; D42-Gal4/UAS-Arf6^CA^* (10.70 ± 0.58 µm^2^). The error bar represents the standard error of the mean (SEM); the statistical analysis was done using one-way ANOVA followed by post-hoc Tukey’s test. *p<0.05, **p<0.01, ***p<0.001; ns, not significant. **(L)** Histogram showing maximum inter-bouton diameter from muscle 4 NMJ at A2 hemisegment in control animals (0.80 ± 0.04 µm), *dAsap^K23/B52^* (1.68 ± 0.07 µm), *actin5C-Gal4, dAsap^K23/B52^; UAS-Arf6^DN^/+* (0.98 ± 0.04 µm), *dAsap^K23/B52^; D42-Gal4/UAS-Arf6^DN^* (0.85 ± 0.04 µm), *dAsap^K23/B52^; mef2-Gal4/UAS-Arf6^DN^* (1.50 ± 0.04 µm), *actin5C-Gal4, dAsap^K23/B52^; UAS-Arf6^CA^/+* (0.87 ± 0.05 µm), *dAsap^K23/B52^; D42-Gal4/UAS-Arf6^CA^* (1.33 ± 0.07 µm). The error bar represents the standard error of the mean (SEM); the statistical analysis was done using one-way ANOVA followed by post-hoc Tukey’s test. *p<0.05, **p<0.01, ***p<0.001; ns, not significant.

To further dissect the epistatic interaction between dAsap and Arf6, we first checked if futsch-positive loops were restored by expressing *Arf6^DN^* in *dAsap* mutants. Consistent with this idea, expressing a dominant negative form of Arf6 in motor neurons (*dAsap^K23/B52^; UAS-Arf6^DN^/D42-Gal4*) restored the futsch-positive microtubule loops in mutants (control, 32.59 ± 1.72 %; *dAsap^K23/B52^*, 17.88 ± 1.09 %, *p*<0.0001; *dAsap^K23/B52^; UAS-Arf6^DN^/D42-Gal4*, 26.33 ± 1.71 %, *p*=0.0003) (Figure 8A-C and 8G). This data indicates the regulation of microtubule-based cytoskeleton by Arf6 effector ASAP in *Drosophila*. Similarly, overexpressing GDP-locked *Arf6^DN^* in motor neurons also normalizes increased active zone number in *dAsap^K23/B52^* mutants (control, 519.2 ± 37.44; *dAsap^K23/B52^*, 1053 ± 107.4, *p*=0.0008; *dAsap^K23/B52^; UAS-Arf6^DN^/D42-Gal4*, 439.5 ± 41.86, *p*=0.0003) (Figure 8D-F and 8H).

**Figure 8.**
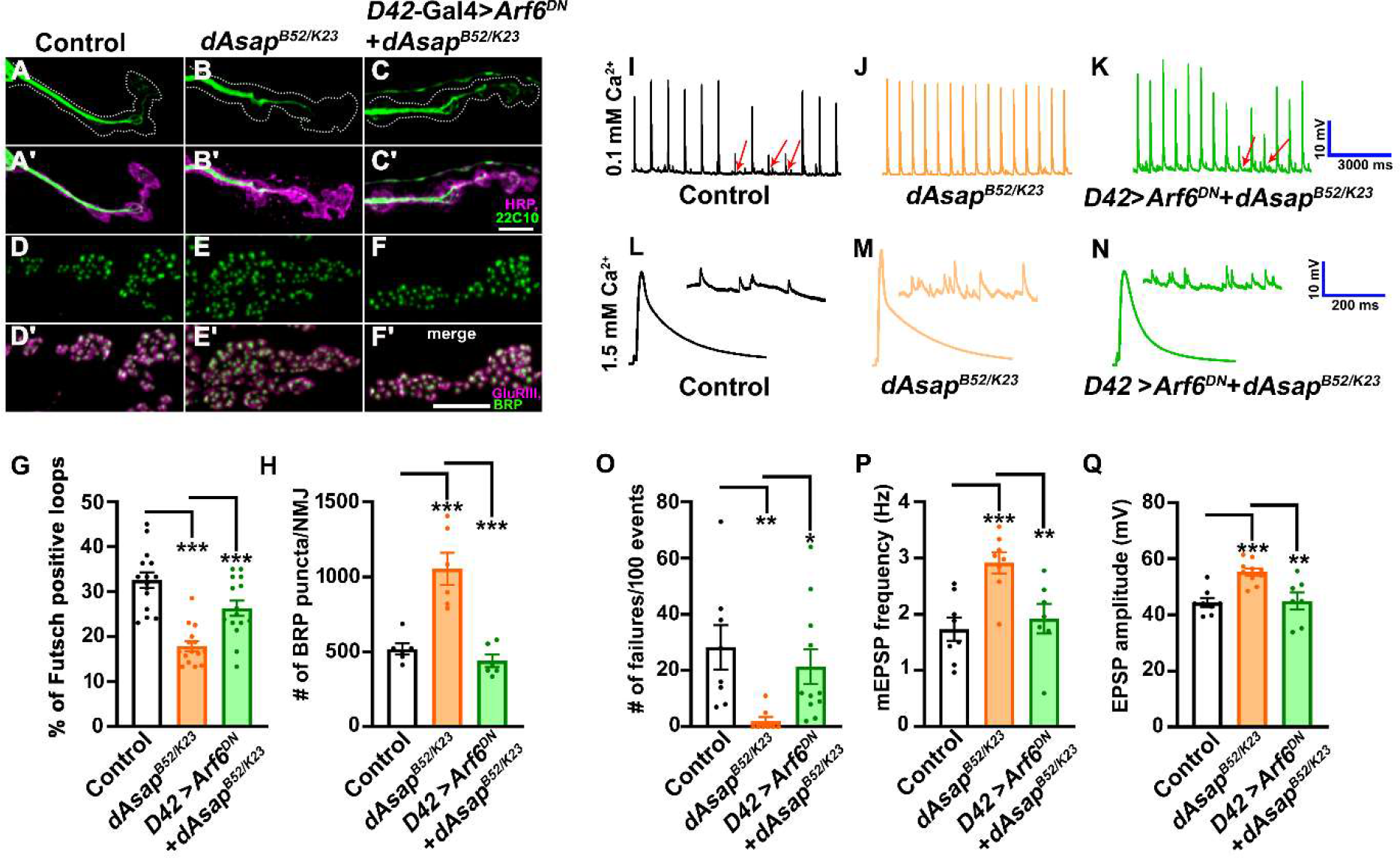
Expression of Arf6^DN^ in motor neurons rescues functional defects of *dAsap* mutants. **(A-C**′) Confocal images of NMJ synapses at muscle 4 of A2 hemisegment showing the futsch loops in (A, A′) control, (B, B′) *dAsap^K23/B52^*, (C, C′) *dAsap^K23/B52^*; *D42-Gal4/UAS-Arf6^DN^* double immunolabeled with 22C10 (green) and a neuronal membrane marker, HRP (magenta). A dominant-negative form of Arf6 expressed in *dAsap* mutant background rescues the reduced futsch loops. Scale bar in C′ (for A-C′) represent 4 μm. **(D-F**′) Confocal images of NMJ synapses at muscle 4 of A2 hemisegment showing active zone density in (A, A′) control, (B, B′) *dAsap^K23/B52^*, (C, C′) *dAsap^K23/B52^*; *D42-Gal4/UAS-Arf6^DN^* double immunolabeled with antibodies against active zones marker Brp (green) and GluRIII (magenta). Note that motor neuron expression of a dominant-negative form of Arf6 in *dAsap* mutant rescue the active zone density. Scale bar in F′ (for D-F′) represent 2.5 μm. **(G)** Histogram showing the percentage of futsch positive loops from muscle 4 NMJ at A2 hemisegment in control animals (32.59 ± 1.721 %), *dAsap^K23/B52^* (17.88 ± 1.099 %), *dAsap^K23/B52^; D42-Gal4/UAS-Arf6^DN^* (26.33 ± 1.712 %). The error bar represents the standard error of the mean (SEM); the statistical analysis was done using one-way ANOVA followed by post-hoc Tukey’s test. *p<0.05, **p<0.01, ***p<0.001; ns, not significant. **(H)** Histogram showing the number of Brp punctae per NMJ from muscle 4 NMJ at A2 hemisegment in control (519.2 ± 37.44), *dAsap^K23/B52^* (1053 ± 107.4), *dAsap^K23/B52^; D42-Gal4/UAS-Arf6^DN^* (439.5 ± 41.86). The error bar represents the standard error of the mean (SEM); the statistical analysis was done using one-way ANOVA followed by post-hoc Tukey’s test. *p<0.05, **p<0.01, ***p<0.001; ns, not significant. **(I-K)** Representative traces of action potential firing at 0.1mM extracellular Ca^2+^ in (I) control, (J) *dAsap^K23/B52^*, (I) *dAsap^B52/K23^; D42-Gal4/UAS-Arf6^DN^* larvae. **(L-N)** Representative traces of mEPSPs and EPSPs at 1.5 mM Ca^2+^ in (L) control, (M) *dAsap^K23/B52^*, *dAsap^B52/K23^; D42-Gal4/UAS-Arf6^DN^* larvae. **(O)** Histogram showing percentage of failure in control animals (28.25 ± 7.90 %), *dAsap^K23/B52^* (2.0 ± 1.42 %), *dAsap^B52/K23^; D42-Gal4/UAS-Arf6^DN^* (21.36 ± 6.16 %). *p<0.05, **p<0.01. The error bar represents the standard error of the mean (SEM); the statistical analysis was done using one-way ANOVA followed by post-hoc Tukey’s multiple-comparison test. **(P)** Histogram showing mEJP frequency from muscle 6/7 NMJ at A2 hemisegment in control animals (1.73 ± 0.20 Hz), *dAsap^K23/B52^* (2.91 ± 0.18 Hz), *dAsap^B52/K23^; D42-Gal4/UAS-Arf6^DN^* (1.92 ± 0.26 Hz). **p<0.01, ***p<0.001. The error bar represents the standard error of the mean (SEM); the statistical analysis was done using one-way ANOVA followed by post-hoc Tukey’s multiple-comparison test. **(Q)** Histogram showing EPSP amplitude from muscle 6/7 NMJ at A2 hemisegment in control animals (44.45 ± 1.59 mV), *dAsap^K23/B52^* (55.28 ± 1.29 mV), *dAsap^B52/K23^; D42-Gal4/UAS-Arf6^DN^* (44.95 ± 3.03 mV). **p<0.01, ***p<0.001. The error bar represents the standard error of the mean (SEM); the statistical analysis was done using one-way ANOVA followed by post-hoc Tukey’s multiple-comparison test.

Next, to understand the inter-relationship between dAsap and Arf6 in the context of synaptic function, we overexpressed *Arf6^DN^*in *dAsap^B52/K23^*mutant combinations. Interestingly, we found that the reduced synaptic failure in *dAsap* mutants was partially restored to the control level (*dAsap^K23/B52^*, 2.0 ± 1.42 %; *dAsap^K23/B52^; UAS-Arf6^DN^/D42-Gal4*, 21.36 ± 6.16 %, *p*=0.017) (Figure 8I-Q and 8O). Moreover, the increased miniature frequency and evoked activity were fully restored by expressing a dominant negative *Arf6* transgene in *dAsap^K23/B52^* mutant combination (mEPSP frequency: *dAsap^K23/B52^*, 2.91 ± 0.18 Hz; *dAsap^K23/B52^; UAS-Arf6^DN^/D42-Gal4*,1.92 ± 0.26 Hz, *p*=0.008) and (EPSP amplitude: *dAsap^K23/B52^*, 55.28 ± 1.29 mV; *dAsap^K23/B52^; UAS-Arf6^DN^/D42-Gal4*, 44.95 ± 3.03 mV, *p*=0.003) (Figure 8L-N and 8P-Q). Taken together, these data indicate that dAsap regulates synaptic function through an Arf6-dependent pathway at the NMJs.

## Discussion

We identified the BAR-domain protein dAsap as one of the regulators of NMJ morphogenesis in a small-scale RNAi-mediated reverse genetic screen (Mallik *et al*., 2017). The presence of Arf GAP domain in dAsap prompted us to investigate the underlying signaling mechanism by which dAsap regulates synaptic morphogenesis and function. Our results highlight three key findings: a) dAsap-dependent Arf6 regulation is required in neurons to promote NMJ growth; b) *dAsap* mutants show perturbed microtubule cytoskeletal organization at the NMJs, and c) dAsap modulates synaptic activity by regulating active zone number and calcium release from intracellular stores at the nerve terminals through a mechanism that involves Arf6-signaling (Figure 9).

**Figure 9.**
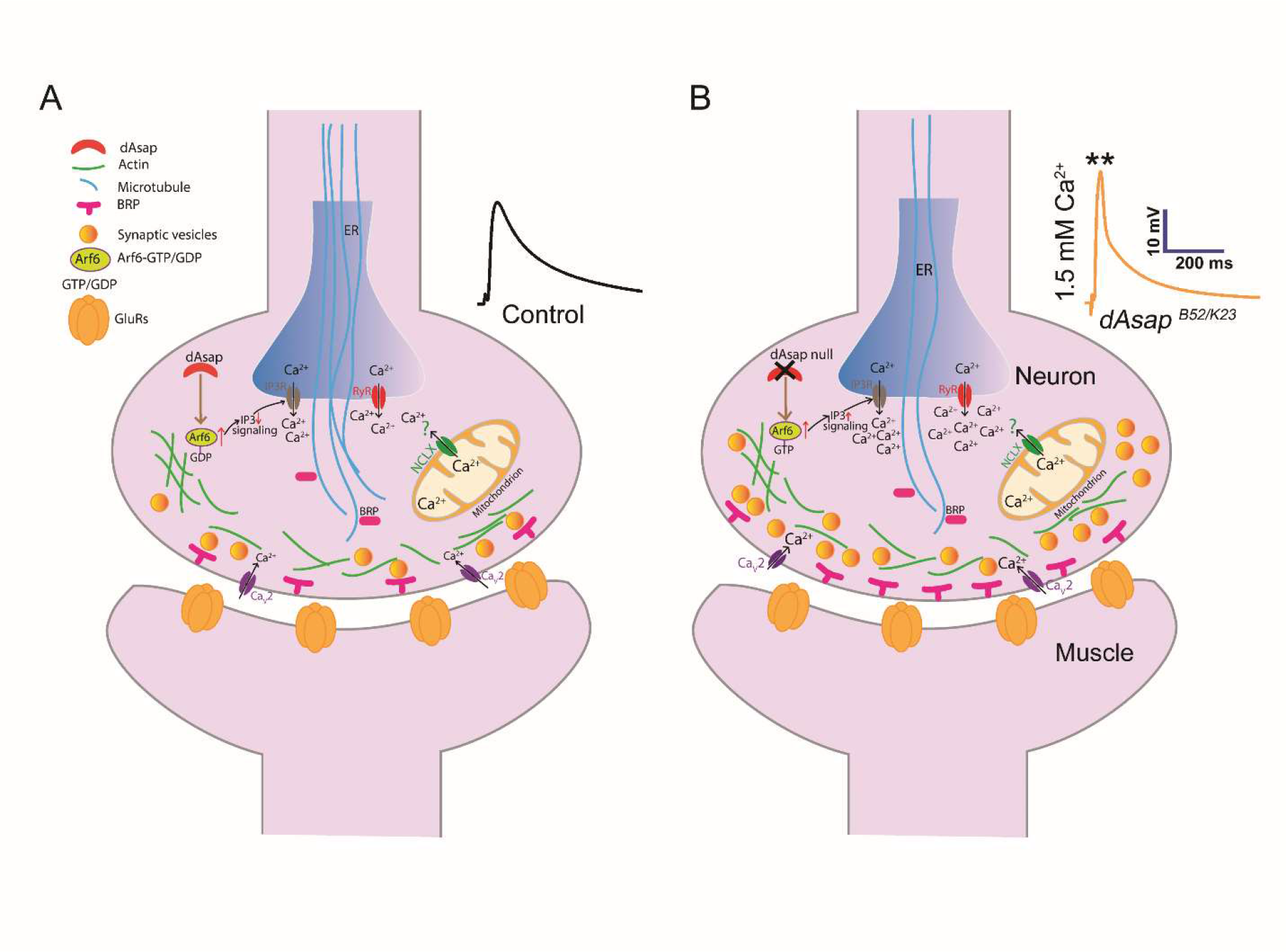
Model depicting regulation of NMJ morphogenesis and function through dAsap-mediated Arf6 signaling in *Drosophila*. Model depicting regulation of NMJ morphogenesis through dAsap-mediated Arf6 signaling in *Drosophila*. Loss of *dAsap* affects the organization of the microtubules at the presynapse, resulting in altered NMJ morphology and increased active zone density at the nerve terminal. The robust increase in evoked neurotransmission in *dAsap* mutants could be due to increased active zone numbers and ER-mediated calcium release that contributes to more vesicle fusion at the synapse. In the wild type, Ca_v_2-type channels maintain normal synapse function; however, in *dAsap* mutants, it might be that elevated Arf6-GTP promotes IP3-mediated calcium release from ER that drives more vesicle fusion at the terminals, which needs further investigation. Besides, further testing is required to dissect if mitochondrial calcium release via the Calx/NCLX channel might contribute to vesicle fusion and hence a robust increase in evoked response in the *dAsap* mutants.

### dAsap regulates NMJ organization through Arf6-dependent signaling

Our studies support the requirement of dAsap in regulating NMJ morphogenesis. The loss-of-function mutants of *dAsap* show impaired synaptic morphology with reduced bouton number and increased bouton size. The boutons seem to be fused with increased inter-bouton diameter. Moreover, the *dAsap* mutant showed disrupted microtubule cytoskeleton loops at synaptic terminals. The tissue-specific rescue experiments reveal that dAsap is required explicitly in neurons to regulate the NMJ structure and microtubule loops.

Previous reports have implicated that mammalian ASAP1 acts as a GAP for Arf proteins (Brown *et al*., 1998). Like its mammalian ortholog, dAsap also has a conserved ArfGAP domain. Our biochemical studies established that dAsap is a GAP for Arf1 and Arf6. Interestingly, dAsap shows more GAP activity towards Arf1 than Arf6, as measured by quantifying the amount of released inorganic phosphate using the Malachite Green based-colorimetric assay. This result implied that Arf1 and Arf6 are substrates for dAsap for GTP hydrolysis, and dAsap could regulate their activities in the neurons during NMJ morphogenesis. However, we did not get any viable larvae for Arf1^DN^ when combined with *dAsap^B52/K23^*; compared to Arf6^DN^ mutant combination. Hence, our genetic data suggest that while Arf1-GTP is the major substrate for dAsap under *in vitro* conditions, it is primarily the Arf6-GTP that is regulated by dAsap in the neurons for the synaptic organization. The Arf family proteins regulate various cellular processes during neuronal development. For instance, Arf1 interacts with PICK1, a BAR domain protein, to modulate the Arp2/3-dependent actin polymerization in mouse primary neurons (Rocca *et al*., 2013). Although we did not observe any significant changes in actin-based moesin-positive GFP puncta at the mutant NMJs; the epistatic interaction with the actin regulatory proteins might answer the involvement of the actin regulator pathway in *dAsap* mutants.

Moreover, several studies have demonstrated that Arf6 is crucial for neurite growth, axon elongation, dendrite development, and synaptic vesicle endocytosis (Hernandez-Deviez *et al*., 2004; Krauss *et al*., 2003; Tagliatti *et al*., 2016). Genetic studies in *Drosophila* suggest that dAsap regulates the activity of Arf1 and Arf6 during cleavage furrow formation and ommatidia patterning in the pupal eye, respectively (Johnson *et al*., 2011; Rodrigues *et al*., 2016). These studies indicate that the role of dAsap could be tissue-specific regulation of Arf proteins. In line with these observations, we found that Arf6^DN^ expression in *dAsap* mutant neurons restores aberrant synapse morphology and defective microtubule cytoskeleton at the NMJ.

*Drosophila* mutants defective in the microtubule organization also exhibit abnormal NMJ growth and function (Lepicard *et al*., 2014; Roos *et al*., 2000). Moreover, defects in the microtubule cytoskeleton at the synapses lead to decreased active zone number and density at the NMJ (Lepicard *et al*., 2014). In contrast, *dAsap* mutants display increased Brp number and density at the terminals. Since expressing a GDP-locked form of Arf6 in the *dAsap* mutant background could rescue the defective microtubule cytoskeleton, one might expect it would also restore the Brp number/density. As expected, neuronal expression of Arf6^DN^ in the *dAsap* mutant background restored the Brp number and density at the terminals. While the mechanism by which dAsap-Arf6 mediated neuronal signaling modulates microtubule organization and Brp number needs further investigation, it is likely that other proteins containing ArfGAP domain along with regulation of other Arfs may work together to regulate synapse organization (Miura *et al*., 2016).

### dAsap modulates robust neurotransmission by inducing active zone density and intracellular calcium release pathway

*dAsap* mutants can sustain synaptic vesicle fusion under low extracellular calcium resulting in significantly fewer synaptic failures. Moreover, loss of *dAsap* results in increased mEJP frequency and EJP amplitude at the NMJ. These electrophysiological defects in *dAsap* mutants seem to arise due to: a) the availability of more intracellular calcium for synaptic vesicle release and b) increased active zone number. In support of this hypothesis, we found a significant increase in the number and density of Brp-punctae at the *dAsap* mutant NMJs. Thus, increased active zone number in *dAsap* mutants could be one of the reasons for increased evoked responses at the terminals.

Studies have revealed the involvement of Arf6-GTP in the regulation of phospholipase C-inositol-3-phosphate (PLC-IP3)-mediated intra-acrosomal calcium release (Pelletan *et al*., 2015). Similar signaling cascades seem to be involved in calcium regulation by intracellular stores and potentiation of synaptic transmission at the neuromuscular synapses. (He et al., 2000; Yang et al., 2001). We speculated that the elevated level of Arf6-GTP might stimulate PLC-IP3-mediated calcium release from intracellular stores in *dAsap* mutants, which could potentiate vesicle fusion. Moreover, the structural analysis of ASAP3 indicates the involvement of ASAP3-ArfGAP domain in complex with Arf6 revealing a conserved arginine residue at 469 positions and a calcium-binding site at the complex interface (Ismail *et al*., 2010). Consistent with this idea, our sequence alignment analysis of dAsap with ASAP3 suggests arginine residue conservation similar to ASAP3 (Figure S3). These analyses indicated that ArfGAP and calcium might be crucial in dAsap-associated GTP hydrolysis, suggesting crosstalk between dAsap, Arf6, and calcium signaling.

Recent studies have shown that IP3-directed signaling stimulates calcium release from IP3R and RyR receptors of the endoplasmic reticulum and is crucial for the fusion of synaptic vesicles during chronic maintenance of homeostasis at the NMJs (James et al., 2019). Moreover, the endoplasmic reticulum facilitates the exchange of calcium and phospholipids by forming dynamic contacts, thus regulating calcium handling in the cells (Paradis et al., 2022). Another independent study suggests that mitochondria control intracellular calcium levels by regulating Na^+^/Ca^2+^ Calx/NCLX exchanger activity and serve as an essential factor in handling the calcium ions in the cells (Palty et al., 2010). Moreover, mitochondrial NCLX exchanger activity in *dAsap* mutants could be one of the potential reasons contributing to calcium levels which need further investigation. These studies support a model where intracellular calcium regulation through Arf6 may regulate synaptic functions.

Altogether, we provide considerable evidence supporting the regulation of intracellular calcium ions at nerve terminals by dAsap-Arf6-dependent signaling. First, blocking IP3 receptors (IP3R) using IP3-sponge and Ryanodine receptors (RyR) using Dantrolene in *dAsap* mutants rescued the EPSP amplitude. Second, consistent with these observations, chelating intracellular calcium-ion using BAPTA significantly restored mEPSP frequency and the EPSP amplitude in the *dAsap* mutants. Third, *dAsap* mutants show lower evoked activity failure rates, indicating a high probability of release due to calcium release from intracellular stores. Expressing a dominant negative form of Arf6 in animals lacking d*Asap* could restore the synaptic failures to wild-type levels. Thus, our studies involving pharmacological and genetic manipulations revealed that dAsap*-*Arf6-GTP regulates intracellular calcium release from the ER through IP3R and RyR-mediated calcium release pathways at the nerve terminals. Notably, this is the first *in vivo* study identifying the Arf6 effector dAsap as a molecule regulating robust neurotransmission at the *Drosophila* NMJs. Overall, our studies support a model in which dAsap-mediated Arf6-dependent signaling regulates a conserved ER calcium handling pathway that modulates the synaptic functions in *Drosophila*.

## Material and Methods

### Drosophila Stocks

Flies were raised and maintained under non-crowded conditions at 25⁰C in standard cornmeal medium (80g/L cornflour, 40g/L dextrose, 20g/L sucrose, 18g/L agar, 15g/L yeast extract, 4%(v/v) propionic acid, 0.06% (v/v) ortho-phosphoric acid and 0.07% methyl-4-hydroxy benzoate/Tego). All the RNAi crosses were set up at 29⁰C unless stated otherwise. All the crosses and rescue experiments were conducted at 25⁰C. *W^1118^* was used as control unless stated otherwise. The Gal4 drivers used in this study were motor neuron-specific *D42-Gal4* (BL-8816), ubiquitous driver *actin5C-Gal4* (BL-25374), and muscle-specific *mef2-Gal4* (BL-50742). *UAS-dAsap-GFP* (BL-65849) was a kind gift from Tony J.C. Harris (University of Toronto, Canada) (Shao et al., 2010). *UAS-ip3-sponge.m30* has been previously described (Usui-Aoki et al., 2005). *UAS-Arf6^DN^* (T27N) (Chen et al., 2003) and *UAS-Arf6^CA^* (Q67L) were obtained from Richa Rikhy (IISER Pune, India). *UAS-Arf1^DN^* (T31N) and *UAS-Arf1^CA^* (Q71L) (Dottermusch-Heidel et al., 2012) were provided by Raghu Padinjat (National Centre for Biological Sciences, India). *UAS-CalX RNAi* (BL-28306) was obtained from Bloomington *Drosophila* Stock Center.

### Generation of *dAsap* loss-of-function mutants and transgene

To generate the loss-of-function mutants of *dAsap*, two sets of gRNAs were designed for the *dAsap* genomic region. The first set of gRNAs was designed at the 2^nd^ and 12^th^ exon, while the second set of gRNAs was designed at the 1^st^ and 12^th^ exon using CRISPR Optimal Target Finder online tool. Primers used to clone these gRNAs into the pCFD4 vector were: gRNA1FP, 5′-TATATAGGAAAGATATCCGGGTGAACTTCGAGATTGGAGTTCGACCGGGGTTTTAGAGCT AGAAATAGCAAG-3′, and gRNA1RP, 5′-ATTTTAACTTGCTATTTCTAGCTCTAAAACCAATAATGCCCTCCATCCAGCGACGTTAAAT TGAAAATAGGTC-3′; gRNA2FP, 5′-TATATAGGAAAGATATCCGGGTGAACTTCGCCCGACTGCCGCCACACGATGTTTTAGAGC TAGAAATAGCAAG-3′, and gRNA2RP: 5′-ATTTTAACTTGCTATTTCTAGCTCTAAAACCGCAATCGTAGAGAGCTCGGCGACGTTAAAT TGAAAATAGGTC-3′. The gRNAs were cloned into a dual gRNA pCFD4 vector having a BbsI restriction site using Gibson Assembly Kit (New England Biolabs (UK) Ltd) following the manufacturer’s guidelines. The pCFD4 vector containing *dAsap* gRNAs was injected into *Drosophila* embryos to generate the transgene. Next, the transgenic flies containing the dAsap gRNAs were crossed with *Nanos-cas9* (BL-54591) to create the deletion of *dAsap* gene in the germline cells. Following standard genetics, lines were established in F2 generation, and *dAsap* deletion was screened by PCR using primers, 5’-TTGGCTAAGTTTGGGAATGC-3’ and 5’-GAGTTTAAAAGACGCATACACACA-3’. Two null mutants *dAsap^K23^*(4190 bp deletion) and *dAsap^B52^* (4275 bp deletion), were obtained.

To generate *dAsap* transgene, a full-length *dAsap* ORF was amplified from cDNA using the following primers and cloned in Gal4-based expression vector pUASt at KpnI and NotI restriction sites: 5′-CCGCCCGGGGATCAGATCCGCATGCCGCCATCCCTGATTG-3′ and 5′-ATCCACTAGTGGCCTATGCGGCCTTAATCAGGCAGCATATGCAC-3′. The pUASt vector containing the *dAsap* ORF was injected into *Drosophila* embryos to generate the transgene.

### Semiquantitative RT-PCR

The expression of *dAsap* in mutants and rescue was analyzed by semiquantitative RT-PCR. In brief, total RNA was isolated from larval fillets using TRIzol reagent (Invitrogen). Reverse transcription was performed on 1 μg total RNA using Superscript^TM^ II Reverse Transcriptase (Invitrogen) using an oligo-dT primer to make cDNA. The resulting cDNA was used for PCR to analyze the level of *dAsap* transcript using primers: 5’-CAAAGATTCCTCCACCAGGA-3’ and 5’-GATCTAGACCGCCACCAGAG-3’. *rp49* was used as an internal control for PCR. The primer sequence used for *rp49* is 5’-AGATCGTGAAGAAGCGCACC-3’ and 5’-CGATCCGTAACCGATGTTGG-3’.

### Generation and affinity purification of antibody

To generate antibody against dAsap, the N-terminal 413 amino acids of dAsap (amino acids 1-413) were cloned in the pET-28a (+) bacterial expression vector. The His-tagged fusion protein was expressed in BL21 codon+ cells, purified from inclusion bodies, and injected into rabbits (Deshpande Laboratories, Bhopal, India). For affinity purification of the anti-dAsap antibodies, 20 μg of the His-tagged dAsap was resolved on 12% SDS-PAGE, transferred to PVDF membrane, and then incubated with 1 mL of anti-dAsap serum diluted 50 folds in 1X PBS (pH 7.4) overnight at 4°C. The membrane was rinsed five times with 1X PBS (pH 7.4). The bound antibody was eluted by incubating the membrane with 500 μl of elution buffer (100 mM glycine; pH 2.5) for 20 minutes at room temperature. The pH was neutralized immediately by adding 0.1 volume of 1.0 M Tris base, pH 10.

### Western blot analysis

Adult fly heads were homogenized in lysis buffer (50 mM Tris-HCl, pH 6.8, 25 mM KCl, 2 mM EDTA, 0.3 M sucrose, and 2% SDS) at 75°C in a water bath. The protein concentration was quantified using bicinchoninic acid (BCA) Protein assay (Sigma-Aldrich). Then 60 μg of the total protein of each genotype was separated on 8% SDS-PAGE and transferred to the PVDF membrane (Amersham, GE Healthcare Life Sciences). The membrane was blocked in 5% skimmed milk (HIMEDIA) in 1X Tris-buffered saline (TBS) with 0.2% Tween-20 (0.2% TBST) for 1 hour at room temperature and then overnight incubated with primary antibody. After washing with 0.2% TBST, the membrane was incubated with HRP-conjugated secondary antibody (1:10000) for 1 hour at room temperature. The primary antibodies used were rabbit anti-dAsap (1:1000) and mouse anti-Ran (1:3000, BD Biosciences). Signals were detected using the Odyssey imaging system (LI-COR Biosciences).

### Immunohistochemistry and quantification

Wandering third instar larvae were dissected in ice-cold Ca^2+^-free HL3 saline and fixed in either 4% PFA for 30 min or Bouin’s fixative (Sigma Aldrich) for 10 min (Mallik *et al*., 2017; Raut et al., 2017). The fillets were incubated with primary antibodies overnight at 4⁰C, followed by secondary antibodies at room temperature for 90 minutes. Finally, larval fillets were washed with 0.2% PBST and mounted with Fluoromount aqueous mounting medium (Sigma-Aldrich) on a glass slide. The monoclonal antibody anti-cysteine string protein (ab49, 1:100), anti-Futsch (22C10, 1:100), and anti-Bruchpilot (nc82, 1:50) were obtained from Developmental Studies Hybridoma Bank (DSHB). Anti-GluRIII (1:100) antibody was a kind gift from Aaron DiAntonio (Washington University, St. Louis). The fluorophore-conjugated secondary antibody Alexa Fluor 488 or Alexa Fluor 568 (Invitrogen, Thermo Fisher Scientific) was used at 1:800 dilution. Alexa Fluor 488 or Rhodamine conjugated anti-HRP (Jackson ImmunoResearch) were used at 1:800 dilution. Images were captured with a laser scanning confocal microscope (Olympus FV3000 or LSM780, Carl Zeiss) using 40x 1.3 NA or 60x 1.42 NA objectives. The images were processed using Image J (ImageJ, NIH, USA) or Adobe Photoshop software (Adobe Inc., USA).

Muscle 4 and muscle 6/7 of A2 hemisegment were used for NMJ morphology quantification. The CSP-positive structures were counted to quantify the total bouton number. For bouton area quantification, CSP-marked boundaries were used to define one bouton. The average of the five biggest boutons from each NMJ was used to quantify bouton size and area. ‘Fused bouton’ phenotype was analyzed using the maximum inter-bouton diameter, measured on three neighboring inter-bouton regions in each sample, and the average of these three regions was used as one data point for quantification(Migh et al., 2018). For Futsch loop quantification, the third instar larval fillets were immunolabelled with HRP and 22C10 antibodies. The NMJs of muscle four were imaged with 40X objective, and futsch-loop structures that localized with HRP were counted. The total number of Futsch loops was divided by the total number of boutons to calculate the percentage of boutons with loops(Roos *et al*., 2000). For multiple comparisons, one-way ANOVA followed by Post-hoc Tukey’s test was used. GraphPad Prism 8 was used to plot the graph. Error bars in bar graphs represent the standard error of the mean (SEM).

### Protein expression and purification

Full-length Arf1 and Arf6 were cloned for biochemical analysis into pGEX-KG bacterial expression vector. The *Escherichia coli* BL21 codon plus cells expressing GST-Arf1 or GST-Arf6 were lysed, affinity purified on glutathione-Sepharose 4B resin column, and digested with Thrombin. The cleaved proteins were eluted and concentrated for further use. The PZA domain of dAsap was cloned in *Pichia pastoris* expression vector pPICZA at KpnI and SacII sites. The His-tagged dAsap^PZA^ protein was expressed and purified following the manufacturer’s guidelines (Invitrogen, USA).

### Malachite green assay

The malachite green substrate is prepared in the ratio of 2:1:1:2 by mixing malachite green (0.08%, w/v in 6N H_2_SO4), ammonium molybdate (5.7% w/v in 6N HCl), polyvinyl alcohol (2.3%, w/v in water with heating) and water (Chang *et al*., 2008; Rowlands et al., 2004). The assay buffer contains 20 mM HEPES pH 7.5, 150 mM KCl, and 5 mM MgCl_2_. Before starting the reaction, 10 μM of each Arf1 and Arf6 purified protein was exchanged with 1 mM GTP in the presence of 3 mM of DMPC, 5 mM EDTA and 0.1% sodium cholate in the assay buffer for one hour at room temperature. The reaction followed the increasing concentration of dAsap^PZA^ (P-PH, Z-ArfGAP, A-Ank) protein starting from 0, 0.1, 0.2, 0.5, 1, 2, 5, and 10 μM and incubated at room temperature for 30 minutes. Further, 160 μl of malachite green reagent was added to the 100 μl of Arf1 or Arf6 and dAsap^PZA^ protein mix in a 96-well plate. The reaction was quenched with 20μl of 3.4% sodium citrate after 5 minutes and kept at 37⁰C for 30 minutes. Next, the absorbance was recorded at 620 nm, and a standard curve was generated using potassium phosphate for every independent experiment.

### Electrophysiology and pharmacology

Intracellular recordings were performed from muscle 6 of A2 hemisegment as described previously (Choudhury *et al*., 2016; Mallik et al., 2023; Mallik and Frank, 2022; Rikhy *et al*., 2002). Briefly, wandering instar larvae were dissected in HL3 saline containing 1.5 mM Ca^2+^ at room temperature. Miniature excitatory postsynaptic potentials (mEPSPs) were recorded for 60 seconds. Evoked excitatory postsynaptic potentials (EPSP) were recorded from the muscle at 1 Hz nerve stimulation. The EPSP was recorded by delivering depolarizing pulse, and the signal was amplified using Axoclamp 900A. The data was digitized using Digidata 1440A and acquired using pClamp7 software (Axon Instruments, Molecular Devices, USA). Intracellular recording microelectrodes having resistance between 20-30 MΩ, filled with the 3M KCl, were used for all recordings. Data from muscles showing resting membrane potential ranging from −60mV to −70mV were used and analyzed. The quantal content was calculated as the ratio of the average EPSP amplitude to the average mEPSP amplitude for each NMJ. As indicated in the figures, the pharmacological agents were mixed in HL3 saline at the final concentrations. The pharmacological agent Dantrolene was obtained from Tocris Bioscience. Failure analysis was performed at 0.1 mM Ca^2+^ containing HL3. One hundred trials (stimulations) were performed at each NMJ in all the genotypes. The failure rate percentage was obtained by dividing the total number of failures by the total number of trials. The data were analyzed using Mini Analysis (Synaptosoft) and Clampfit software (Molecular Devices, USA).

## Acknowledgements

We thank Dr. Ruth Johnson for sharing DsRed-tagged dAsap transgene and Dr. Tony Harris for shRNA transgenes against dAsap and GFP-tagged-dAsap transgenes. We thank the Bloomington Drosophila Stock Center (BDSC), Vienna Drosophila RNAi Centre (VDRC), and Drosophila Genomics Resource Centre (DGRC) for fly stocks and Developmental Studies Hybridoma Bank (DSHB), the University of Iowa for monoclonal antibodies. We thank Dr. Manish Jaiswal and Dr. Shanker Jha for their helpful comments on the manuscript.

